# Zinc arrests axonal transport and displaces tau, doublecortin, and MAP2C from microtubules

**DOI:** 10.1101/2021.12.02.470968

**Authors:** Taylor F. Minckley, Lyndsie A. Salvagio, Dylan H. Fudge, Kristen Verhey, Steven M. Markus, Yan Qin

## Abstract

Accurate delivery of cargo over long distances through axonal transport requires precise spatiotemporal regulation and relies on microtubule function. Here we discover that Zn^2+^ influx via depolarization inhibits axonal transport. Zn^2+^-mediated inhibition is nonselective for cargo. Elevated Zn^2+^ (IC_50_ ≈ 5-10 nM) reduces both lysosomal and mitochondrial motility in primary rat hippocampal neurons and HeLa cells. We further reveal that Zn^2+^ directly binds to microtubules, inhibiting movement of motor proteins (kinesin and dynein) and promoting detachment of neuronal-specific MAPs (Tau, DCX, and MAP2C). We finally provide a detailed model of microtubule interactions with Tau, DCX, dynein, kinesin, and predict microtubule Zn^2+^ binding sites. Our results reveal that Zn^2+^ acts to inhibit the microtubule binding of tau, DCX, and MAP2C and can directly block the progression of motor proteins on microtubules. Intraneuronal Zn^2+^, therefore, is a critical signal for regulating axonal transport and microtubulebased processes.

## Introduction

Free, labile zinc (referred to herein as Zn^2+^) has been suggested to act as an intraneuronal signaling molecule. Increases in cytoplasmic Zn^2+^ concentrations can occur as a result of influx from the extracellular milieu through several channels historically defined as Ca^2+^ channels^1^. Alternatively, Zn^2+^ can be released from intracellular stores via the transient receptor potential mucolipin 1 (TRPML1) channel^2,3^. Synchronous Zn^2+^ spikes have also been detected to fire along with neuronal excitation^4^. Such elevated Zn^2+^ has long-term transcriptional effects in neurons^5,6^. Bioinformatic studies have suggested that Zn^2+^ can bind 2800 human proteins^7^, and *in vitro* studies have shown that physiological levels of Zn^2+^ inhibits certain enzymes^8–10^. However, it has yet to be determined whether changes in intraneuronal Zn^2+^ can rapidly elicit downstream effects at timescales comparable to Ca^2+^ signaling events. Moreover, if and how Zn^2+^ might directly affect neurophysiological processes is poorly understood.

Microtubule-based trafficking plays critical roles in neurons, which have axonal and dendritic projections that can extend up to ~1 meter in humans. The importance of proper axonal transport and microtubule regulation is evident by the clear correlation between several neurodevelopmental and adult-onset neurodegenerative diseases and mutations in several components of microtubule network and transport machinery^11–15^. These processes are under precise spatiotemporal control for the establishment and maintenance of neuronal function and health. Microtubule-associated proteins (MAPs) are responsible for the stability and regulation of microtubule dynamics and microtubule-based function, including transport. Motor proteins comprise one subfamily of MAPs and include dynein and kinesin which are responsible for retrograde and anterograde trafficking along microtubules, respectively^16,17^. Recent work has established that various MAPs can influence polarized microtubule transport by either inhibiting or promoting motor protein interaction with microtubules^18^. A subset of these motility- and stability-regulating MAPs are upregulated in neurons, and are crucial for neuronal development and function, including MAP-Tau (Tau), doublecortin (DCX), and MAP2C. Together, motor proteins (and their cargo-binding adaptor proteins), MAPs, and the microtubules themselves provide numerous targets for regulation that help the cell accomplish appropriate transport and microtubule regulation.

In this manuscript, we uncover potent roles for Zn^2+^ in the regulation of microtubule-based processes. Neuron depolarization-induced Zn^2+^ influx arrested axonal transport of lysosomes and mitochondria, which was reversed upon Zn^2+^ chelation. Furthermore, physiological levels of Zn^2+^ directly inhibited motility of kinesin via a cargo-adaptor independent mechanism by interacting with microtubules. To determine the potential Zn^2+^ binding sites on microtubules, we investigated the effects of Zn^2+^ on microtubule-decoration of eight MAPs by confocal imaging. Our results show that increases in cytosolic Zn^2+^ induce fast and reversible dissociation of Tau, DCX, and MAP2C from microtubules. We propose that Zn^2+^, via direct interactions with microtubules, acts as a brake for axonal transport and redistributes DCX, MAP2C and Tau in neurons.

## Results

### Zn^2+^ arrests axonal transport of lysosomes and mitochondria

Both Ca^2+^ and Zn^2+^ were reported to inhibit mitochondrial motility in neurons^19–21^. This led us to question whether this inhibition is limited to mitochondria, or if it could affect other organelles such as lysosomes. In order to determine the effects of Zn^2+^ on lysosomal trafficking, we increased intracellular Zn^2+^ concentrations in primary hippocampal neurons expressing LAMP1-mCherry by triggering depolarization-induced extracellular Zn^2+^ influx^22^. Membrane depolarization with 50 mM KCl in the presence of 100 μM ZnCl_2_ led to a significant decrease in the motility of lysosomes (Figure 1A-E, Video S1). Upon treatment with 100 μM of the Zn^2+^ chelator TPEN (N,N,N’,N’-tetrakis(2-pyridinylmethyl)-1,2-ethanediamine), motility immediately recovered to near-baseline levels. Although both retrograde and anterograde lysosomal motility (speed and proportion) were rescued to baseline levels upon addition of TPEN, anterograde motility was slower to recover. Conversely, neuron depolarization with 50 mM KCl and 2 mM CaCl_2_ caused no significant reduction in lysosome motility, and rather, there was an increase in the velocity and proportion of anterograde-moving lysosomes (Figure 1F-I, Video S2).

**Figure 1:**
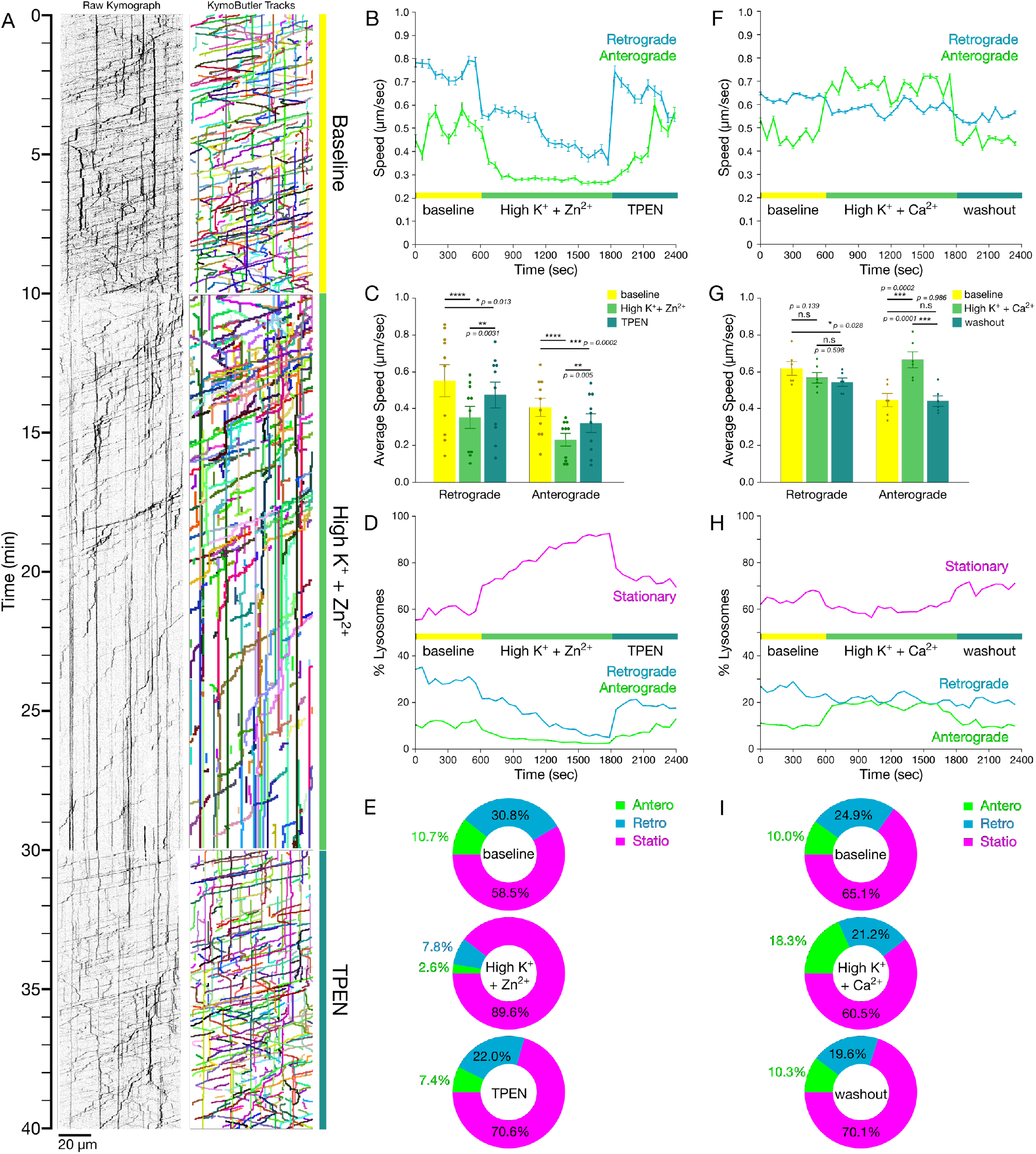
Depolarization-induced influx of Zn^2+^, not Ca^2+^, arrests axonal transport of lysosomes. (A) Representative kymographs (left) and measured tracks (identified using KymoButler; right) of LAMP1-mCherry motility along axons from primary cultured rat hippocampal neurons prior to (“baseline”; top) or after initiation of Zn^2+^ influx induced by 50 mM KCl depolarization (middle), and then following washout and addition of 100 μM TPEN (bottom). (B) Mean instantaneous speed (±s.e.m.) of lysosomes moving retrograde (blue) or anterograde (green) across baseline, Zn^2+^ influx by 50 mM KCl depolarization, and TPEN treatment (60-second binned, representing 10 axons). (C) Mean speed (±S.E.M.) of lysosomes moving in the indicated directions for each condition (n = 10 axons from 6 independent replicates). Least-squares regression with post-hoc Tukey HSD. (D) Proportions of lysosomes moving in the indicated directions across baseline, depolarization, and TPEN treatment (60-second binned, representing 10 axons). (E) Mean proportions of lysosomal motility for each condition. (F) Mean instantaneous speed (±s.e.m.) of lysosomes moving retrograde (blue) or anterograde (green) across baseline, 2 mM Ca^2+^ influx by 50 mM KCl depolarization, and washout (60-second binned, representing 10 axons). (G) Mean speed (±S.E.M.) of lysosomes moving in the indicated directions for each condition (n =10 axons from 6 independent replicates). Least-squares regression with post-hoc Tukey HSD. (H) Proportions of lysosomes moving in the indicated directions across baseline, 2 mM Ca^2+^ with 50 mM KCl, and washout (60-second binned, representing 10 axons). (I) Mean proportions of lysosomal motility for each condition.**** *p* < 0.0001, *** *p* < 0.001, ** *p* < 0.01, * *p* < 0.05, n.s. not significant.

We also observed that Zn^2+^ influx significantly decreased the motility of mitochondria marked by mito-mCherry (Figure S1A-E, Video S3), a phenomenon which has been previously reported ^21^. Upon TPEN addition, we observed a fast rescue of retrograde mitochondrial transport, but a slower recovery of anterograde transport. In addition, the recovered motility with TPEN treatment could be completely inhibited again by treating the cells with 100 μM ZnCl_2_ and 2.5 μM pyrithione (a Zn^2+^ ionophore), indicating a highly reversible mechanism (Figure S1A-H). These results suggest that neuronal Zn^2+^ acts as a reversible brake on the axonal transport of both lysosomes and mitochondria, implicating a universal role of Zn^2+^, distinct from Ca^2+^, in regulating microtubule-based trafficking.

### Zn^2+^ inhibits organellar motility with a nanomolar IC_50_ in HeLa cells and neurons

We wondered whether Zn^2+^ could arrest microtubule-based trafficking in non-neuronal cell types and found that inhibition of organellar motility could also be seen in HeLa cells upon addition of 20 μM ZnCl_2_ and 1.25 μM pyrithione (Figure S2). As with neurons, organellar motility could be rescued with TPEN (Figure S2, Video S4). Copper can act as a competitor for certain Zn^2+^ binding sites on proteins^23^, so we tested whether copper could induce the same inhibition of motility. Surprisingly, we found no inhibition of lysosomal motility in HeLa cells upon addition of 20 μM CuCl_2_ and 2.5 μM pyrithione (Figure S3). We then sought to determine the cytoplasmic concentration of Zn^2+^ that is necessary to inhibit motility. We simultaneously quantitated organellar motility and cytosolic Zn^2+^ concentrations by expressing either LAMP1-mCherry or mito-mCherry, and the genetically encoded Zn^2+^ sensor, GZnP2 (K_d_ = 352 pM for Zn^2+^)^24^ in HeLa cells (Figure 2A). Zn^2+^ was chelated with TPEN to determine the minimum apo-state signal of GZnP2, followed by 100 μM ZnCl_2_ to slowly increase cytosolic Zn^2+^. Cells were then treated with 2.5 μM pyrithione to achieve maximal GZnP2 signal (Figure 2B-C). Organellar motility was measured and plotted against the corresponding cellular Zn^2+^ concentrations (see Methods), generating doseresponse curves for Zn^2+^-dependent inhibition of both lysosomes (Figure 2D & H, IC_50_ = 7.01 ± 2.08 nM) and mitochondria (Figure 2E & H, IC_50_ = 5.50 ± 2.51 nM). To confirm our HeLa cell findings, we simultaneously measured axonal transport of lysosomes and axoplasmic Zn^2+^ concentration in primary hippocampal neurons, and observed a strikingly similar threshold of Zn^2+^ inhibition (Figure 2F-H, IC_50_ = 7.38 ± 2.80 nM).

**Figure 2:**
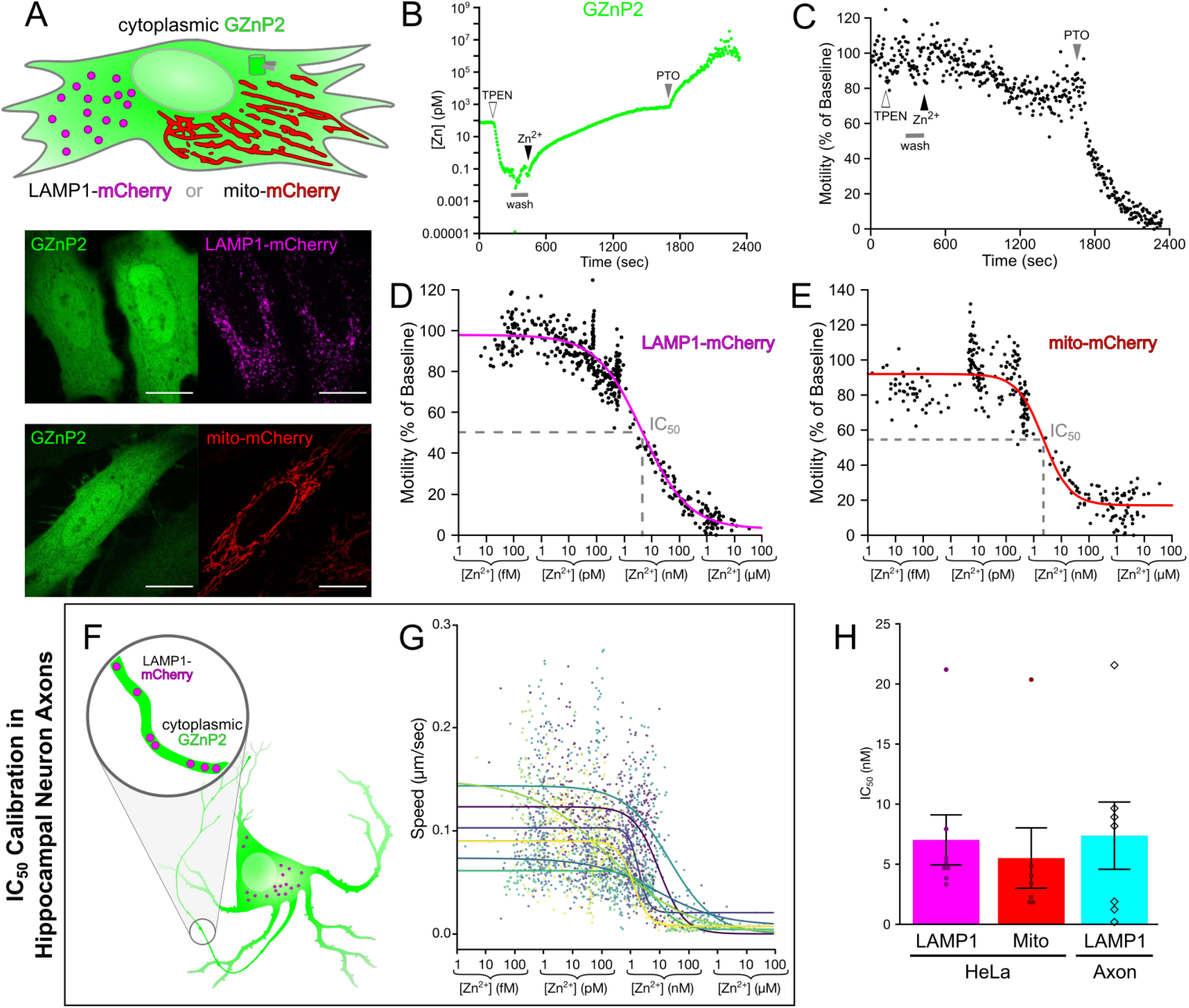
Zn^2+^ inhibits organellar motility with nanomolar IC_50_ in HeLa cells and neurons. (A) Schematic of the IC_50_ assay in HeLa cells (top) showing cytosolic localization of co-transfected GZnP2 sensor and either lysosomal (left, magenta) or mitochondrial (right, red) mCherry. Representative micrographs of HeLa cells (bottom) illustrating the localization of GZnP2 sensor in the cytoplasm, and either LAMP1-mCherry (magenta, top) or mito-mCherry (red, bottom). Scale bar = 20 μm. (B and C) Representative plots of [Zn^2+^] (B; determined using GZnP2; see Methods) and relative lysosomal motility (C; via LAMP1-mCherry) in a HeLa cell treated as indicated (100 μM TPEN at t = 120 sec; a 2 min duration washout at t = 300 sec; 100 μM ZnCl_2_ at t = 420 sec; and 5 μM pyrithione at t = 1680 sec). (D and E) Representative plot of relative motility versus [Zn^2+^] used to calculate IC_50_ values Zn^2+^ inhibition of (D) LAMP1-mCherry or (E) mito-mCherry motility. (F) Schematic of the IC_50_ assay in primary hippocampal neurons showing cytosolic localization of co-transfected GZnP2 sensor (green) and LAMP1-mCherry tagged lysosomes (magenta). (G) Overlaid plots of KymoButler-analyzed lysosome speed versus [Zn^2+^] used to calculate IC_50_ values for Zn^2+^ inhibition of lysosomes in neuron axons. (H) Mean IC_50_ values (±s.e.m.) for LAMP1-mCherry (“LAMP1”) in HeLa cells (magenta; n = 8 cells from 6 individual replicates) or primary rat hippocampal neuron axons (cyan; n = 7 neuron axons from individual replicates) and mito-mCherry in HeLa cells (“Mito”, red; n = 7 cells from 5 individual replicates. All experiments were performed in the absence of extracellular Ca^2+^.

### Zn^2+^ inhibits kinesin motility in a dose dependent manner but does not cause microtubule detachment

Next, we investigated whether Zn^2+^-mediated inhibition is a consequence of direct modulation of motor or cargo-adaptor proteins by employing an inducible cargo trafficking assay. Constitutively active motors are recruited to transport vesicular cargoes in a cargo-adaptor-independent manner in living cells^25^. A green fluorescent protein (mNeonGreen) and rapamycin binding fragment (FRB) are fused to the C-terminus of a constitutively active fragment of the neuron-specific kinesin-1 motor protein (KIF5A(1-560)-mNG-FRB)^17^. Two FK506 binding proteins (2xFKBP) and a red fluorescent protein (mRFP) are fused to the C-terminus of a peroxisomal membrane targeting sequence (PEX3(1-42)-mRFP-2xFKBP). When rapamycin or its analogs (i.e. Zotarolimus) are added to this system, FRB-FKBP heterodimerization is induced and the motor protein and peroxisome are irreversibly tethered (Figure 3A), resulting in peroxisomes being transported from their generally static perinuclear position to the cell periphery^25,26^. With this system, we could directly assess KIF5A motility by measuring peroxisome movement and localization.

**Figure 3:**
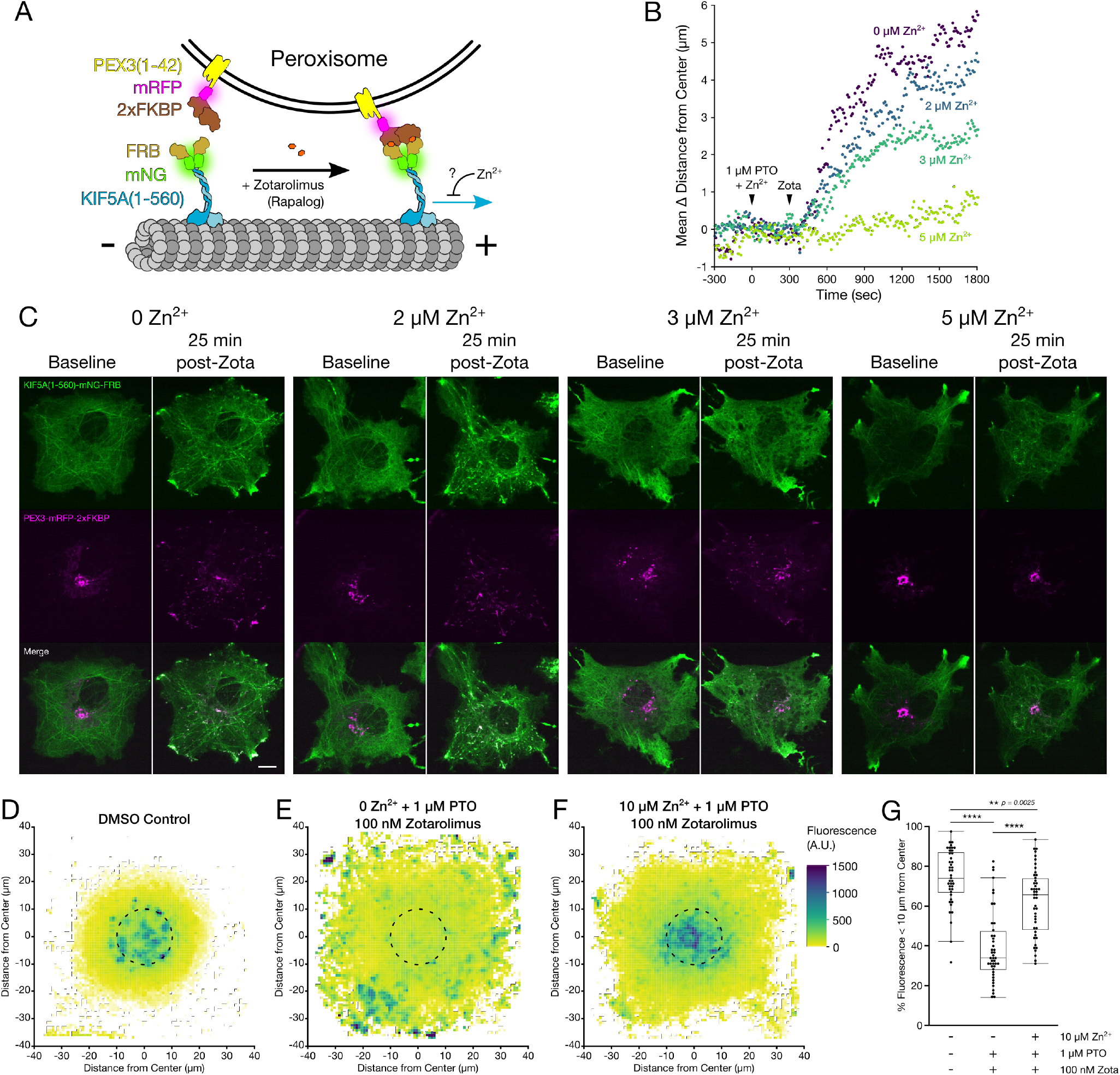
Zn^2+^ inhibits KIF5A movement in situ in a dose dependent manner. (A) Schematic of the components of the peroxisome dispersion assay. (B) Representative plots of mean change in peroxisome distance from the cell center over time, for COS-7 cells expressing KIF5A(1-560)-mNG-FRB and PEX3-mRFP-2xFKBP, treated at 0 sec with 1 μM PTO and various concentrations of Zn^2+^ (none, purple; 2 μM, blue; 3 μM, teal; or 5 μM, lime), and 100 nM Zotarolimus (“Zota”) at 300 sec. (C) Representative micrographs of COS-7 cells expressing KIF5A(1-560)-mNG-FRB (green, top), PEX3-mRFP-2xFKBP (magenta, middle), or merged channels (bottom), at 0 seconds (“baseline”) or 25 minutes after treatment with 100 nM Zotarolimus (“post-Zota”) in the presence of the indicated Zn^2+^ concentrations. Scale bar = 10 μm. (D-F) Fluorescent intensity averages of PEX3-mRFP-2xFKBP in COS-7 cells co-expressing KIF5A(1-560)-mNG-FRB. Cells were imaged and aligned by their geometric centers after the following treatments and fixation: (D) 2 μL DMSO for 5 minutes, then an additional 2 μL DMSO for 25 minutes (n = 42 cells); (E) 1 μM PTO for 5 minutes, then 100 nM Zotarolimus for 25 minutes (n = 44 cells); (F) 10 μM ZnCl_2_ and 1 μM PTO for 5 minutes, then 100 nM Zotarolimus for 25 minutes (n = 46 cells). Black dotted line indicates the area within 10 μm of the center. (G) Box plots and individual points representing the proportion of PEX3-mRFP-2xFKBP fluorescence within 10 μm of the geometric cell center, for the each of the conditions in (D-F). One-way ANOVA with post-hoc Tukey HSD. All experiments were performed in the absence of extracellular Ca^2+^. **** *p* < 0.0001, ** *p* < 0.01.

COS-7 cells were co-transfected with KIF5A(1-560)-mNG-FRB and PEX3-mRFP-2xFKBP^26^. Cytosolic KIF5A(1-560)-mNG-FRB fluorescent signal was visibly recruited to the peroxisomal membranes within seconds of Zotarolimus treatment (Figure 3C) and peroxisomes were rapidly dispersed to the periphery (Figure 3C, Video S5), measured by the average change in distance of peroxisomes from the geometric center of the cell (Figure 3B). Strikingly, when ZnCl_2_ (2, 3, or 5 μM) was added along with pyrithione, peroxisome dispersion was dramatically reduced in a dose-dependent manner (Figure 3B & C, Video S5). We quantified the inhibitory effect on peroxisomal distribution after 25 minutes of Zotarolimus treatment. Radial analysis of fluorescence intensity from the geometric cell center was conducted using a custom image processing macro written for ImageJ, and representative averages of all analyzed cells were generated (Figure 3D-F). Zotarolimus treatment led to a significant dispersion of peroxisomes compared with DMSO controls, which was almost entirely inhibited after Zn^2+^ influx (Figure 3G). Kinesin was still visibly recruited to peroxisomes in the presence of 5 μM Zn^2+^ (Figure 3C), indicating that Zn^2+^ does not interfere with FRB-FKBP heterodimerization.

Using oblique illumination microscopy we observed that KIF5A(1-560)-mNG-FRB was highly enriched at the cell periphery and exhibited linear morphology, which colocalized with mCherry-α-tubulin signal, confirming that KIF5A interacts with and walks along microtubules toward their plus ends (Figure S4A). After 5 minute treatment with 20 μM ZnCl_2_ and 2.5 μM pyrithione, KIF5A(1-560) signal significantly shifted toward the center of the cell and away from the periphery (Figure S4A), quantified by the ratio of edge:center fluorescence intensity (Figure S4C). Pyrithione treatment alone had no effect on the distribution of KIF5A(1-560) (Figure S4B-C). Confocal timelapse imaging revealed redistribution of KIF5A(1-560), but no changes in the distribution of mCherry-α-tubulin (Figure S4D, Video S6), suggesting that Zn^2+^ addition did not cause a massive microtubule catastrophe event. It is important to note that Zn^2+^ did not cause dissociation of KIF5A from microtubules (Figure S4A). Therefore, Zn^2+^ modulates motility by acting on the motor proteins or microtubules.

### Zn^2+^ directly binds microtubules and differentially modulates microtubule-stimulated ATPase activity of kinesin and dynein in vitro

Our live cell imaging data clearly demonstrated that Zn^2+^ inhibits motor protein motility. Previous *in vitro* studies showed that millimolar concentrations of Zn^2+^ assembles purified tubulin into 2-dimensional sheets and macrotubes^27^ and micromolar Zn^2+^ inhibits the ATPase activity of kinesin^28^, though we observed *in situ* effects on motility in the nanomolar range. We therefore used accurately-buffered Zn^2+^ solutions (from the picomolar to micromolar range) to examine the dose-dependent Zn^2+^ effects on microtubule-stimulated ATPase activity of motor proteins *in vitro.* We tested a commercially available human recombinant kinesin heavy chain (KIF5A, #KR01 Cytoskeleton) as well as an artificially dimerized, truncated yeast dynein motor domain fragment, which is sufficient for processive motility^29^ and exhibits high similarity to human dynein.

We tested the microtubule-stimulated ATPase activity of these motors at varying Zn^2+^ concentrations using the EnzChek ATPase assay. Interestingly, we observed that KIF5A ATPase activity increases in response to picomolar levels of Zn^2+^ (peaking around 100 pM), but then is reduced as Zn^2+^ increases into the nanomolar range (Figure 4A, Figure S5A). Alternatively, dynein activity is relatively stable across all picomolar Zn^2+^ concentrations and exhibits a similar decrease as Zn^2+^ increases into the nanomolar and micromolar range (Figure 4B, Figure S5B). Baseline ATPase activity (rate of ATPase per motor domain per second, “ATPase·motor^-1^·sec^-1^”) for both kinesin (~2-5 ATPase·motor^-1^·sec^-1^) and dynein (~8-9 ATPase·motor^-1^·sec^-1^) is similar to the previously-reported values (Figure S5) ^30–33^. Finally, we observed that there was little to no reduction of ATPase activity of either kinesin or dynein in the presence of 500 μM Ca^2+^ (Figure 4C).

**Figure 4:**
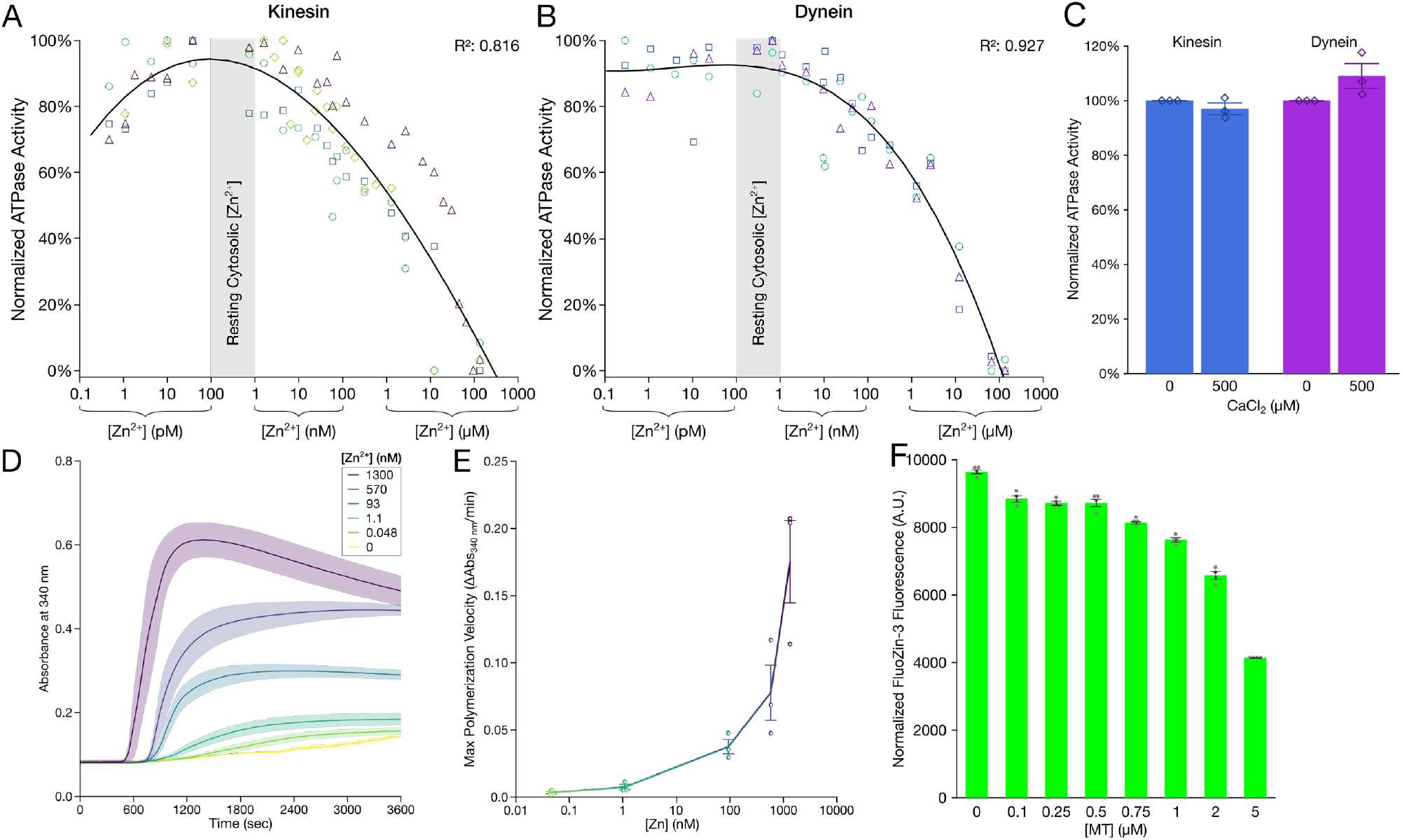
Zn^2+^ can bind microtubules and inhibits motor protein activity in vitro. (A-B) Microtubule-stimulated ATPase activity of (A) purified recombinant human kinesin (KIF5a), and (B) a minimally processive, artificially dimerized yeast dynein fragment (GST-dynein_331_) across a range of precisely-buffered Zn^2+^ concentrations (see Methods), normalized to the maximum and minimum ATPase activity within each individual replicate, replicates denoted by separate colors and marker shapes (kinesin, n = 4 individual replicates; dynein, n = 3 individual replicates). Each assay was performed in the presence of 2 μM microtubules. Black line shows cumulative data fitted to cubic model with log transformation of [Zn^2+^] concentration, with corresponding R^2^ values shown in the top right corner. (C) Mean normalized microtubule-stimulated ATPase activity (±S.E.M.) for a minimally processive, artificially dimerized yeast dynein fragment (GST-dynein_331_, purple) and recombinant human kinesin (KIF5A, blue) in the presence of 500 μM CaCl_2_ (n = 3 replicates each). (D-E) Traditional turbidity assay of microtubule polymerization across a range of precisely-buffered Zn^2+^ concentrations (see Methods). (D) Mean absorbance (±S.E.M.) at 340 nm and (E) maximum polymerization velocity for each Zn^2+^ concentration (n = 3 replicates per Zn^2+^ concentration). (F) Mean (±S.E.M.) *in vitro* fluorescent intensity (488 nm excitation, 525 nm emission) for solutions containing 1 μM FluoZin-3, 1 μM ZnCl_2_, 1 mM DTT, and varied concentrations of preformed, taxol-stabilized porcine microtubules (MT) (n = 4 individual replicates). **** *p* < 0.0001, n.s. not significant.

Given that both kinesin and dynein ATPase activity can be inhibited by Zn^2+^ *in vitro,* we reasoned that Zn^2+^ might directly act on microtubules to inhibit motor protein motility. Previous studies have demonstrated potential Zn^2+^ binding sites on tubulin^27,34^. The small molecule Zn^2+^ sensor FluoZin-3 (K_d_ = 15 nM) was used to determine whether pre-polymerized canonical microtubules could chelate labile Zn^2+^ out of solution containing the sensor and Zn^2+^. As microtubule concentration increased, FluoZin-3 fluorescence intensity decreased (Figure 4D), indicating that microtubules can bind Zn^2+^. Further, we sought to observe whether nanomolar concentrations of Zn^2+^ could influence microtubule polymerization. Strikingly, Zn^2+^ increased the rate and extent of microtubule polymerization in a dose-dependent manner (Figure 4E-F). Together, these *in vitro* assays suggest that Zn^2+^ can directly bind to microtubules in cells.

### Zn^2+^ promotes the dissociation of tau, doublecortin, and MAP2C from microtubules in situ

Given the finding that decoration of microtubules with Zn^2+^ inhibits the progression of motor proteins, which are one type of a vast array of MAPs, Zn^2+^ binding may affect microtubule interactions of other MAPs. To test this, we increased intracellular Zn^2+^ in COS-7 cells with 20 μM ZnCl_2_ and 2.5 μM pyrithione and examined the microtubule decoration of eight different MAPs. Strikingly, Zn^2+^ caused dissociation of EGFP-Tau, EGFP-DCX, and EGFP-MAP2C from microtubules within 5 minutes as measured by “tubeness” analysis (Figure 5A-G, Video S7), without affecting microtubule morphology (Figure S6A). In addition, microtubule association of EGFP-Tau, EGFP-DCX, and EGFP-MAP2C could be rapidly rescued with Zn^2+^ chelation using 100 μM TPEN (Figure 5A-G, Video S7), suggesting a highly reversible mechanism as seen in the motility experiments. No decrease in microtubule association was seen with pyrithione treatment alone, nor with influx of Ca^2+^ (Figure 5E-G). In addition, we tested whether copper had similar effects, as tau has been shown to bind copper^35^. Interestingly, 20 μM CuCl_2_ and 2.5 μM pyrithione had no effect on EGFP-DCX or EGFP-MAP2C, while both 20 μM CuCl_2_ and 100 μM CuCl_2_ and 2.5 μM pyrithione caused a slight but statistically insignificant dissociation of EGFP-tau from microtubules in some cells (Figure 5F). Zn^2+^ had no effects on the microtubule interaction of GFP-MAP1B, mEmerald-MAP4, EGFP-MAP7, EGFP-MAP9, or EGFP-p150glued (Figure 5H). We next examined if endogenous axonal tau was dissociated from microtubules by Zn^2+^. Primary rat hippocampal neurons were treated for 5 minutes with 50 mM KCl with or without the presence of 100 μM ZnCl_2_, and immediately fixed with −20 °C methanol to simultaneously permeabilize cells and preserve microtubule morphology^36^. Immunofluorescence with anti-Tau antibodies showed clear decoration along axons in neurons treated with KCl alone, but Zn^2+^ caused a drastic reduction in axonal Tau localization (Figure 5I-J, Figure S7).

**Figure 5:**
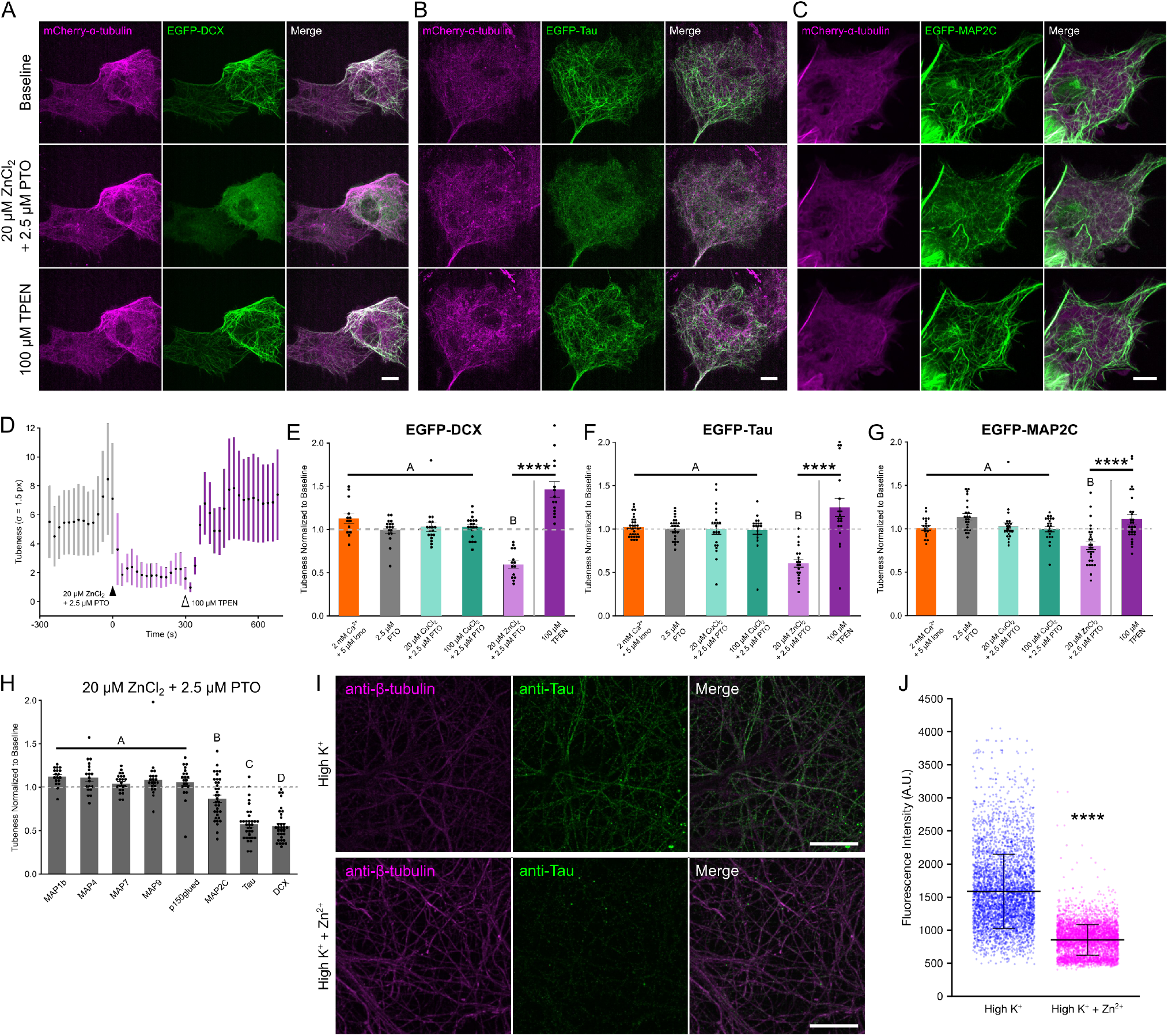
Zn^2+^ promotes reversible detachment of DCX, tau, and MAP2C from microtubules in situ. (A-C) Representative micrographs of COS-7 cells expressing mCherry-α-tubulin (magenta, left) and either (A) EGFP-DCX, (B) EGFP-Tau, or (C) EGFP-MAP2C (green, center), and merged channels (right), before treatment (“baseline”, top), after 5-minute treatment with 20 μM ZnCl_2_ and 2.5 μM PTO (middle), or after subsequent treatment with 100 μM TPEN (bottom). Scale bar = 10 μm. (D) Representative time trace of microtubule morphology of EGFP-DCX in COS-7 cells as measured by “Tubeness” analysis (see Methods and Supplementary Figure 6). Cells were treated with 20 μM ZnCl_2_ at 0 sec and 100 μM TPEN at 300 sec. (E-G) Mean tubeness (±S.E.M.) of (E) EGFP-DCX, (F) EGFP-Tau, or (G) EGFP-MAP2C normalized to baseline (gray dashed line), for the indicated treatments. Letters indicate statistically different groups (see Supplementary Tables 1–3). One-way ANOVA with post-hoc Tukey HSD. Zn^2+^ chelation rescue (100 μM TPEN) was measured with paired t-test. (H) Mean tubeness (±S.E.M.) of the indicated microtubule-associates proteins in COS-7 cells treated with 20 μM ZnCl_2_ and 2.5 μM PTO for 5 minutes, was normalized to baseline. Letters indicate statistically different groups (see Supplementary Table 4). One-way ANOVA with post-hoc Tukey HSD. (I) Representative immunofluorescence micrographs of methanol-fixed primary rat hippocampal neurons depolarized with 50 mM KCl in the absence (top) or presence (bottom) of 100 μM ZnCl_2_. Fixed cells were incubated with anti-β-tubulin monoclonal antibodies (magenta, left), anti-Tau polyclonal antibodies (green, center), and channels are shown merged (right). Scale bar = 10 μm. (J) Fluorescence intensity of AlexaFluor 488 secondary antibody-labeled Tau decorating linear tubulin-positive regions across a 1 mm^2^ area of neurons depolarized with 50 mM KCl in the absence (blue) or presence (magenta) of 100 μM ZnCl_2_ (see Figure S7) (50 mM KCl: n = 4330 linear segments from 2 biological replicates; 50 mM KCl and 100 μM ZnCl_2_: n = 4820 linear segments from 2 biological replicates). Unpaired t-test assuming unequal variances. All experiments were performed in the absence of extracellular Ca^2+^, except where indicated. **** *p* < 0.0001.

### An integrated atlas of microtubule-MAP interactions and predicted Zn^2+^ binding sites

Previous cryo-EM work has provided near-atomic models of microtubules^37^ and their interaction with MAPs, including Tau^38^, DCX^39^, kinesin^40^, and dynein^41^. In order to simultaneously visualize where these MAPs bind on the microtubule lattice, we used the Protein-Ligand Interaction Profiler (PLIP)^42^ to identify the amino acid residues on α- and β-tubulin which interacted with each of these MAPs (Figure 6A). All these MAPs bind in unique areas along the microtubule surface, with most binding along the “ridge” of the protofilaments. Though these MAPs interacted with microtubules in distinct areas, all four MAPs had some interaction with helix-12^43^ of α-tubulin. We wondered whether there are overlapping sites among these four MAPs and Zn^2+^ that can account for the Zn^2+^-induced dissociation of Tau and DCX and inhibition of motor protein motility. In order to explore this, we performed bioinformatic analysis on the 3D protein structure of microtubules using the MIB Server^44^, and rendered the 3D model of microtubules to visualize these binding sites (Figure 6B). Surprisingly, there were a number of domains on the outer surface of the microtubule with high prediction scores for Zn^2+^ binding, specifically on helix-12. There was a significant overlap between the predicted Zn^2+^ binding sites and the MAP-interacting residues (Figure 6A-B), as well as residues within 3Å of these sites (Figure S8). We identified a cluster of residues with both high Zn^2+^ binding scores and significant MAP interactions, including H192, E196, E417, E420, E423, D424. One site involving H192, E420, and D424 is particularly favorable for Zn^2+^ binding to form a tetrahedral geometry within 2-3 Å distance to a Zn^2+^ ion (Figure 6Bii). One water molecule or E196, which directly interacts with DCX (Figure 6Ai) and is only 5.3 Å from this Zn^2+^ site, may act as the fourth Zn^2+^ coordination site. Tau has been found to sterically hinder motor protein binding to microtubules^45^. Given the location of Zn^2+^ binding sites on microtubule surface, Zn^2+^ could compete with MAPs for the same amino acid binding sites, or induce a conformational change in these MAP-binding regions of microtubules. This could cause the dissociation of Tau and DCX, and Zn^2+^ could act similarly to tau to block motor proteins.

**Figure 6:**
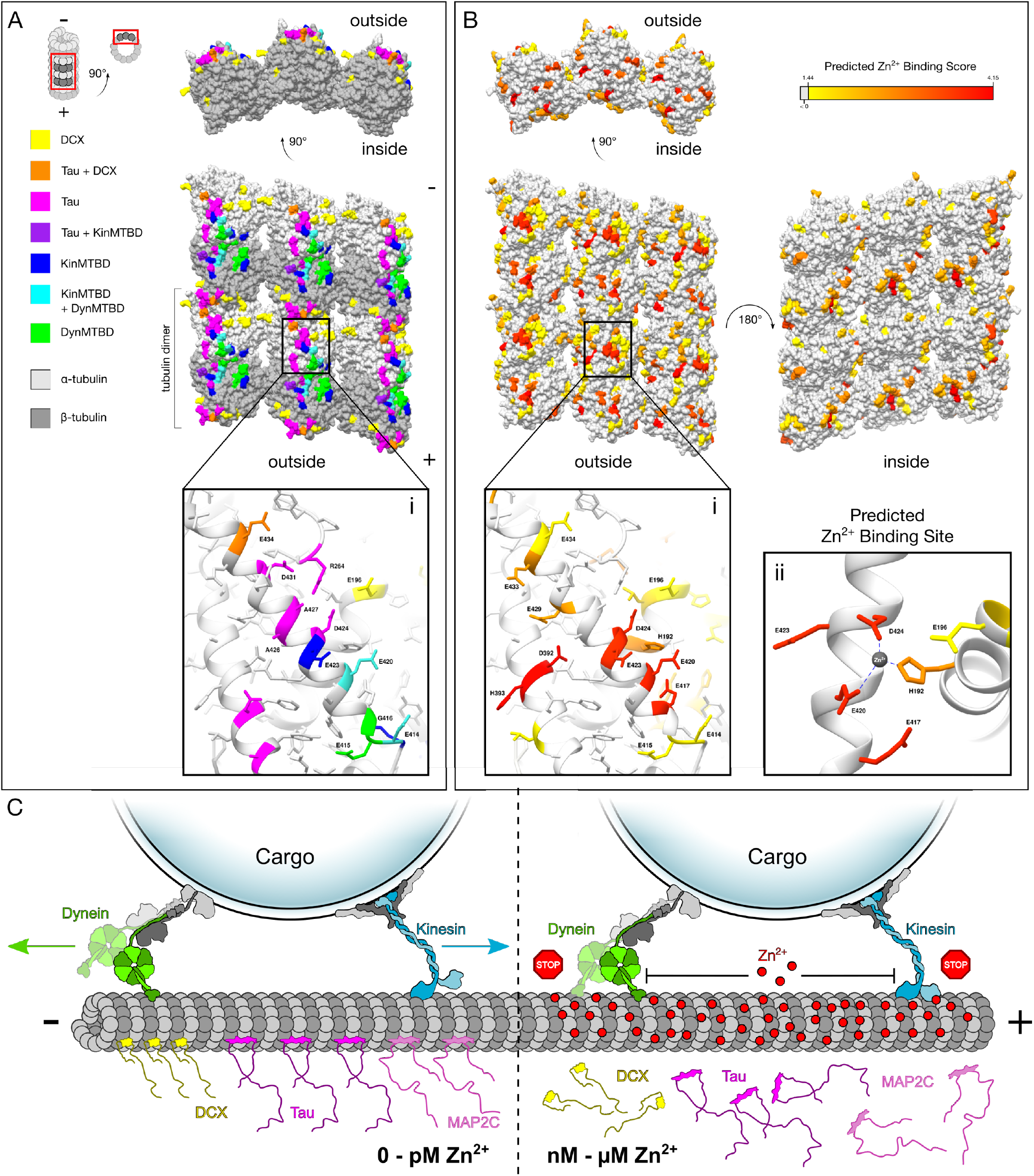
Analysis of MAP and Zn^2+^ binding sites on microtubules, and mechanistic model of microtubule regulation. (A-B) 3D protein model of cryo-EM microtubule reconstructions (PDB ID: 6EVY), with 6 tubulin dimers shown across 3 protofilaments (red boxes from inset schematic). (A) Amino acids of α-tubulin (light gray) and β-tubulin (dark gray) are color-coded to represent which MAPs directly interact at that residue as calculated by the Protein-Ligand Interaction Profiler (PLIP) from cryo-EM reconstruction models for DCX (PDB ID: 6REV), Tau (PDB ID: 6CVN), kinesin microtubule binding domain (“KinMTBD”, PDB ID: 3J8X), and dynein microtubule binding domain (“DynMTBD”, PDB ID: 6RZB). (B) Tubulin amino acids are color coded to represent predicted Zn^2+^-binding sites (red = high score, yellow = moderate score, light gray = low score) as calculated by the MIB Server (see methods)^44^. Insets (i) in each black box show the amino acids on or near helix 12 of alpha tubulin that interact directly with the color-coded MAPs (A) or that are predicted Zn^2+^ binding residues (B). (ii) Predicted Zn^2+^ binding site between H192, E420, and D424. Dotted lines represent 2-3 Å distances between the predicted Zn^2+^ binding residues and the Zn^2+^ ion (gray). (C) Proposed model of Zn^2+^-dependent motility inhibition and MAP detachment upon cytosolic Zn^2+^ increase. Increased cytosolic Zn^2+^ (red circles) in the nanomolar to micromolar range can bind to microtubules and inhibit motor protein (kinesin and dynein) progression, pausing organellar transport. Zn^2+^ binding to microtubules can also cause detachment of DCX, Tau, and MAP2C.

## Discussion

Long-distance axonal transport requires coordinated and precise regulation to ensure delivery of appropriate cargo to its requisite destination. Our results reveal that Ca^2+^ does not inhibit lysosomal motility, but Zn^2+^ can inhibit axonal transport of both mitochondria and lysosomes. Such inhibition was detected when Zn^2+^ concentration was increased to low nanomolar range (IC_50_ = 5 nM, Figure 2), which is within the range of previous estimates of the physiological fluctuations in cytosolic Zn^2+^ concentration^46^. We further clarify that Zn^2+^-dependent inhibition of organelles is due to a direct effect on motor protein activity (Figure 3). Our *in vitro* work suggests that Zn^2+^ ions directly bind to sites along microtubules (Figure 4D-F, Figure 6C). Furthermore, we demonstrate that nanomolar concentrations of Zn^2+^ were able to reduce the ATPase activity of both kinesin and dynein *in vitro* (Figure 4A-B). The slight discrepancy in inhibitory concentration between *in vitro* and *in situ* (high nanomolar vs low nanomolar) could be due to a combinatorial effect of Zn^2+^ on MAPs and motor proteins. We discovered that elevated levels of Zn^2+^ can selectively detach tau, DCX, and MAP2C from microtubules, without disrupting other MAPs (Figure 5). Further, this Zn^2+^-dependent detachment occurs within seconds, and is completely reversible. This highlights the importance of Zn^2+^ homeostasis on second-to minute timescales, as changes in Zn^2+^ concentration can acutely modulate microtubule-based processes. Interestingly, all three Zn^2+^-regulated MAPs have established roles in neuronal development and morphological determination of axons and dendrites^47,48^, further suggesting a crucial regulatory role of Zn^2+^ in neurons. Since dynein, kinesin, and three MAPs are all affected by Zn^2+^, one or more Zn^2+^ binding sites may exist on the microtubule at a shared binding region, or at a site that exhibits allosteric effects on the interaction between MAPs and microtubules (Figure 6).

As neurons rely on the well-coordinated transport of various axonal cargoes with a high degree of spatial and temporal precision, these findings provide an important new role for Zn^2+^ and the consequences of its imbalance in the development and progression of neuronal disease. Given the different responses of dynein and kinesin to low concentrations of Zn^2+^, changes in local Zn^2+^ concentration can potentially favor either anterograde or retrograde trafficking events. This could play an important role in the balance of motor protein “tug-of-war” for cargoes which are decorated with both kinesin and dynein. Long-term changes in Zn^2+^ concentrations may also signal a cell-wide change in motility, wherein motility in the anterograde *(i.e.*, high picomolar to low nanomolar Zn^2+^), retrograde *(i.e.,* low picomolar Zn^2+^), or neither (*i.e.*, > 10 nM Zn^2+^) direction is favored. Slight changes in cytosolic Zn^2+^ concentration could result in the perturbation of the fine-tuned regulation of axonal cargo transport between the soma and axonal growth cones or synaptic terminals, and chronic Zn^2+^ dyshomeostasis may contribute to wide-scale transport defects. Defects in axonal transport can lead to several adult onset neurodegenerative diseases, including Alzheimer’s disease, Huntington’s disease, amyotrophic lateral sclerosis (ALS), spinal muscular atrophy with lower extremity dominance (SMA-LED), and Parkinson’s disease^11^. Unsurprisingly, mutations in several components of the transport machinery *(e.g.,* dynein, cargo adaptors, LIS1, kinesin) have been linked to both neurodevelopmental and neurodegenerative diseases^12,13^. Neurons are especially susceptible to dysfunctions in transport regulation due to their highly elongated and extensively branched morphology. Thus, in order to understand the mechanism underlying the establishment and maintenance of neuronal health and develop strategies to prevent or reduce the severity of disease, it is imperative that we understand the mechanisms that can affect and regulate axonal transport. Our findings reveal insight into a new mode of regulation of axonal transport.

Given the relatively high concentration of Zn^2+^ in neurons^22^ and high dynamics of neuronal Zn^2+^ signals^2,4^, our work reveals a new regulation mechanism of microtubule-based processes and highlights Zn^2+^ as an ionic signal for proper neurodevelopment and neuronal function. Local increases in cytosolic Zn^2+^ can result from influx through VGCCs in the soma or axon terminals ^22,49,50^, or NMDA-dependent Zn^2+^ influx at synapses^22,51^. These local Zn^2+^ signals may cause organelles to stop or pause in order to deliver their cargo, such as Shank proteins, to the postsynaptic regions during synaptogenesis and synapse maturation^52^. Here, we show that even short-term (5 minute) elevations in Zn^2+^ are able to drastically reduce endogenous binding of Tau to microtubules (Figure 5I-J).

Such local or global elevation of Zn^2+^ may also lead to spatiotemporal control of DCX in growing neurites, as well as proper redistribution of tau and MAP2C to axons and dendrites, respectively. Global Zn^2+^ increases have also been shown during ischemia^53,54^ and in Alzheimer’s disease patients^55,56^. Abnormal Zn^2+^ elevation under pathological conditions could disrupt proper localization of MAPs and importantly promote tau detachment from microtubules, thus increasing the chance for tau aggregates to form in the cytosol. In fact, Zn^2+^ has been shown to directly influence tau aggregation^57–59^. Our work supports existing hypotheses that Zn^2+^ imbalance may be a precursor to Alzheimer’s disease and other tauopathies^34,60,61^, while also revealing a possible mechanism of Zn^2+^ homeostasis for regulation of axonal transport and neuronal MAP function. From a broader perspective, this study corroborates a long-speculated signaling role of intracellular Zn^2+^ with the discovery that Zn^2+^ can directly regulate microtubule-based organellar trafficking, which is essential and required for diverse cellular processes in all eukaryotic cell types throughout the evolutionary spectrum.

## Acknowledgements

This work was funded by the NIH (R01NS110590 to Y.Q.; R01GM118492 and R35GM139483 to S.M.M; R35GM131744 to K.V.)

## Author contributions

T.F.M. contributed experimental design, live and fixed-cell fluorescence imaging, *in vitro* assays, data analysis and interpretation, figure generation, and wrote the manuscript. L.A.S. contributed live cell imaging, data analysis and interpretation, and figure generation. D.H.F. contributed experimental design, live cell imaging, data analysis and interpretation. K.V. contributed design and construct of peroxisome dispersion assay based on FRB-FKBP heterodimerization. S.M.M. contributed protein purification, experimental design, data interpretation, and wrote the manuscript. Y.Q. supervised the project and contributed experimental design, data interpretation, and wrote the manuscript.

## Materials and Methods

### Animal care

Pregnant Sprague Dawley rats were purchased from Charles River (strain: 400). Animal treatment and maintenance were performed by the University of Denver Animal Facility (AAALAC accredited). All experimental procedures using animals were approved by the Institutional Animal Care and Use Committee (IACUC) of the University of Denver.

### Plasmid generation

Expression plasmids were constructed using traditional molecular biological techniques and were all verified by sequencing. To construct LAMP1-mCherry, mCherry was PCR amplified from mCherry-Rab7a-7, LAMP1 was PCR amplified from LAMP1-RFP, and both were ligated into pcDNA3.1+. mCherry-Rab7a-7 was a gift from Michael Davidson (Addgene plasmid #55127; http://n2t.net/addgene:55127; RRID:Addgene_55127). Lamp1-RFP^62^ was a gift from Walther Mothes (Addgene plasmid # 1817; http://n2t.net/addgene:1817; RRID:Addgene_1817). mCh-alpha-tubulin^63^ was a gift from Gia Voeltz (Addgene plasmid # 49149; http://n2t.net/addgene:49149; RRID:Addgene_49149). EGFP-DCX^64^ was a gift from Joseph Gleeson (Addgene plasmid # 32852; http://n2t.net/addgene:32852; RRID:Addgene_32852). pRK5-EGFP-Tau^65^ was a gift from Karen Ashe (Addgene plasmid # 46904; http://n2t.net/addgene:46904; RRID:Addgene_46904). pEGFP-p150Glued^66^ was a gift from David Stephens (Addgene plasmid # 36154; http://n2t.net/addgene:36154; RRID:Addgene_36154). GFP-MAP1B^67^ was a gift from Phillip Gordon-Weeks (Addgene plasmid # 44396; http://n2t.net/addgene:44396; RRID:Addgene_44396). pEGFP-Map7-fulllength^68^ was a gift from Edgar Gomes (Addgene plasmid # 46076; http://n2t.net/addgene:46076; RRID:Addgene_46076). mEmerald-MAP4-C-10^69^ was a gift from Michael Davidson (Addgene plasmid # 54152; http://n2t.net/addgene:54152; RRID:Addgene_54152). MAP9 in pCR-XL-TOPO (#BC146864) was purchased from Transomic Technologies, and subcloned into pcDNA with an N-terminal EGFP tag. MAP2C in pENTR223-1 (#BC172263) was purchased from Transomic Technologies, and subcloned into pcDNA with an N-terminal EGFP tag.

### Primary rat hippocampal neuron culture and transfection

Primary hippocampal neurons were prepared from rat embryos at embryonic day 18 (E18) in dissection medium containing HBSS, 1 M HEPES buffer (pH 7.3) and 50 μg/mL gentamycin. The hippocampi were minced and treated with 1000 U/mL papain added to the dissection medium and dissociated by trituration in 1 mg/mL DNase I. Cells were plated on 1 mg/mL poly-L-lysine-coated 10 mm round glass coverslips at a density of 20,000–25,000 cells/coverslip per dish in neuron plating medium (MEM supplemented with glucose and 5% FBS). After checking that cells were adhered, neuron plating medium was replaced with Neurobasal medium (Thermo Fisher Scientific, Waltham, MA) supplemented with 0.3X GlutaMAX (Thermo Fisher Scientific) and 1X B-27 (Thermo Fisher Scientific). Cultures were maintained at 37 °C, 5% CO_2_. Neurons were transfected between 5 and 10 days *in vitro* (DIV), using the Lipofectamine 3000 transfection kit (Thermo Fisher Scientific) in 500 μL Opti-MEM. The reagent-DNA mixture was incubated for a minimum of 25 min at room temperature before direct addition to the neuron imaging dishes. Before adding reagent-DNA, 1 mL media was removed from each imaging dish and syringe filtered with an equal volume of fresh neuron culture media (50:50 media). After incubation at 37 °C for 4 hours, neurons were washed three times with 1 mL prewarmed neuron culture media. Then, 2 mL of the 50:50 media was added to the neuron imaging dishes and they were incubated at 37 °C until imaging.

### Drug treatment and live cell imaging of neurons, and analysis of axonal motility

Live cell imaging of neurons was performed two or more days after transfection. Neurons were washed and imaged in HEPES-Buffered Hanks Balanced Salt Solution (HHBSS) lacking Ca^2+^ and Zn^2+^. For experiments in which Zn^2+^ influx was induced by depolarization, neurons were treated with Ca^2+^-free HHBSS supplemented with 50 mM KCl and 100 μM ZnCl_2_ during the acquisition period (20 minutes in duration). For experiments in which chelation followed depolarization-induced Zn^2+^ influx, neurons were washed 3 times with Ca^2+^-free HHBSS, then treated with 100 μM TPEN and immediately imaged. For experiments in which Zn^2+^ influx was induced with pyrithione treatment, neurons were washed 3 times with Ca^2+^-free HHBSS and then treated with 100 μM ZnCl_2_ and 2.5-5 μM pyrithione.

All live neuron imaging was performed on an inverted Nikon/Solamere CSUX1 spinning disc confocal microscope equipped with a 40x 1.4 NA oil immersion objective, and a variable 1.0x or 1.5x OptiVar tube lens. Axonal motility imaging was performed in a LiveCell imaging chamber kept at 37°C, 5% CO_2_, and 80% humidity. The microscope was controlled with MicroManager software and image analysis was performed using Fiji^70^ (see below for details on image analysis). To image lysosomal and mitochondrial motility (via LAMP1-mCherry or mito-mCherry, respectively), fluorescence images were acquired every 0.5–2.5 seconds, with 50-100 millisecond exposures with a 561 nm laser at 1-5 mW power.

To analyze organellar axon motility, image contrast of each movie was enhanced by first employing a bandpass filter to remove frequencies larger than 7 px and less than 2 px. Fiji’s built in “Background subtraction” was then run using a 5 px rolling ball radius. Image stacks were then stabilized (if needed) using the Manual Drift Correction plugin for Fiji. Kymographs were generated using the Kymolyzer plugin^71^, and background kymograph noise was removed using a custom code (ImageJ Macro Language). These kymographs were then analyzed using the KymoButler^72^ plug-in for Mathematica (Wolfram Research, Inc.). Instantaneous velocities measured by the KymoButler plug-in were extracted using custom code (Wolfram Language) in order to get a more accurate description of motility, as the density of bidirectionally moving lysosomes reduced the accuracy of computer-generated track averages. Axons were manually traced back to their soma to confirm orientation, which was used to classify retrograde or anterograde motility. KymoButler-measured tracks with durations less than 4 seconds were also removed entirely from analysis to eliminate the measurement of kymograph noise artifacts.

### Non-neuronal cell culture and transfection

HeLa cells and COS-7 cells (African Green Monkey fibroblast-like cell line) were maintained in high glucose Dulbecco’s Modified Eagle Medium (DMEM) supplemented with 10% fetal bovine serum (FBS). COS-7 cells were grown in 100 U/mL Penicillin G and 100 μg/mL Streptomycin, on poly-D-lysine coated coverslips. All cells were maintained at 37°C and 5% CO_2_. HeLa and COS-7 cell transfections were performed with polyethyleneimine (PEI; dissolved to 1 mg/ml in water at pH 7.2, with long-term storage at −20°C and short-term storage at 4°C). For each transfection, 3-6 μL of PEI and 1-1.25 μg of DNA were mixed in 250 μL Opti-MEM, incubated for > 25 min at room temperature, then added to one imaging dish with cells (~40-50% confluent). Imaging dishes were incubated at 37°C until imaged as described above.

### Zn^2+^ concentration measurement using genetically encoded fluorescent probes

For simultaneous analysis of Zn^2+^ concentrations and organellar motility, GZnP2 was co-transfected into HeLa cells or primary rat hippocampal neurons with either mito-mCherry or LAMP1-mCherry. After acquiring images to determine baseline Zn^2+^ concentration and motility, 100 μM TPEN was added to chelate Zn^2+^. Cells were then washed 3 times to remove TPEN, treated with 100 μM ZnCl_2_, then treated with 2.5 μM pyrithione (as indicated on respective plots). GZnP2 fluorescence was acquired by excitation with 488 nm and collecting 525 nm emission. GZnP2 fluorescence intensity values were converted to Zn^2+^ concentrations using the equation,

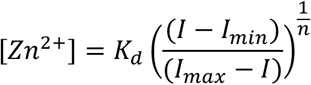

where K_d_ is the Zn^2+^ equilibrium dissociation constant, *n* is the Hill coefficient, *I* is background-subtracted fluorescence intensity, and *I_min_* and *I_max_* are minimum and maximum fluorescence intensities, respectively. K_d_ and Hill coefficients were experimentally determined using purified GZnP2, which revealed K_d_ = 352 pM, and n = 0.49^24^. Data points that yielded [Zn^2+^] > 100 μM (no more than 3 data points out of 300+ total data points per cell) were removed from further analysis since theoretically cytosolic [Zn^2+^] shouldn’t exceed the concentration of the exogenous buffer. Normalized motility values (for HeLa cells, see below) or direct speeds (for neuron axons, see KymoButler methods above) were plotted against the calculated Zn^2+^ concentrations, and fit to a sigmoidal curve using JMP Pro 15.2.0 or KaleidaGraph (Synergy Software), which was used to calculate the Zn^2+^ IC_50_ concentration. Data were fit to sigmoidal curves using a standard dose-response curve,

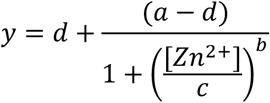

where *a* is the upper asymptote, *b* is the growth rate, *c* is the IC_50_, and *d* is the lower asymptote.

### Organellar motility analysis in HeLa cells

The Total Motility plugin^73^ for Fiji was used to determine the relative motility of organellar structures. Images were binarized by automatic thresholding, then processed by the Total Motility plugin. Motility data were normalized to baseline using the equation,

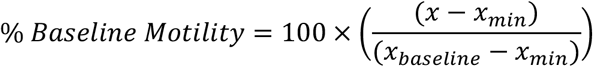

where *x* equals the raw motility data output from the Total Motility plugin, *x_min_* equals the minimum raw motility data point, and *x_baseline_* equals the average motility value for all baseline timepoints. The raw motility data output from the Total Motility plugin is the proportion of thresholded binary pixels that changed from one frame to the next, as a percentage of the total number of positive (white, non-background) pixels in that frame. Images of LAMP1-mCherry or mito-mCherry were acquired every 5 seconds with a 50–100 millisecond exposure of 561 nm laser excitation at 1–5 mW power. The raw motility output was normalized to average baseline motility for each cell. Extreme motility changes that were clearly due to observable stage shift (*i.e.* during drug addition) were removed from further analysis.

Lysosomal or mitochondrial motility was measured using the “Total Motility” plugin^73^ for ImageJ, which calculates the percentage of organellar movements by measuring the area of shifted mitochondria (or lysosomes) out of the total mitochondrial (or lysosomal) area for each acquisition time point in each cell.

### Peroxisome dispersion assays and analysis in COS-7 cells

For rapalog-induced peroxisome dispersion assays, COS-7 cells were co-transfected with PEX3-mRFP-2xFKBP and either KIF5A(1-560)-mNG-FRB or KIF5C(1-559)-mNG-FRB, and imaged within 24 hours as longer transfection resulted in altered localization of the PEX3 construct and cell morphology. Peroxisome dispersion assays were imaged on the Nikon/Solamere CSUX1 spinning disc confocal microscope as described above. For timelapse experiments, cells were imaged for a 5-minute baseline, then 1 μM pyrithione and varying Zn^2+^ concentrations (as indicated) for 5 minutes. Finally, 100 nM Zotarolimus was added to induce heterodimerization, and cells were imaged for 25 minutes. Images were acquired every 10 seconds with a 50-200 ms exposure for both 488 nm and 561 nm laser excitation at 10 mW and 5 mW power, respectively. Custom code (ImageJ Macro Language) was used to detect PEX3-mRFP maxima and quantify average distance of peroxisomes from the center of the cell.

As a non-dispersed peroxisome control, COS-7 cells were treated with 2 μL DMSO for 5 min (vehicle control for pyrithione), then another 2 μL DMSO for 25 minutes (vehicle control for Zotarolimus). As a positive control, COS-7 cells were treated with 1 μL PTO for 5 minutes, 100 nM Zotarolimus for 25 minutes. Finally, to test the effect of Zn^2+^, COS-7 cells were treated with 10 μM Zn^2+^ and 1 μM PTO for 5 minutes, 100 nM Zotarolimus for 25 minutes. After treatment, cells were fixed with 4% paraformaldehyde in 0.1 M PBS with 4% sucrose at 4 °C for 10 minutes. Fixation was quenched with 0.1 M glycine for 5 minutes, then cells were washed with PBS and imaged. Custom code (ImageJ Macro Language) was used to measure fluorescence intensity of PEX3-mRFP as a function of distance from the geometric cell center.

### Oblique illumination imaging of motor proteins and microtubules in COS-7 cells

COS-7 cells were co-transfected with KIF5A(1-560)-mNG-FRB and mCherry-α-tubulin, and imaged within 48 hours. Images were taken using a ONI Nanoimager with oblique illumination (55.1° illumination angle), 1-2% 488 nm laser power, 6-8% 561 nm laser power, and 25 ms to 100 ms exposure times. High-frequency images for in situ kinesin motility (Figure S7A) were taken at 40 Hz.

### Yeast dynein complex protein purification

Purification of the artificially dimerized (by glutathione-S-transferase, GST) minimal dynein motor fragment (GST-dynein^MOTOR^-HaloTag) was performed as previously described^74^ Briefly, yeast cultures were grown in YPA supplemented with 2% glucose (for the intact dynein complex) or 2% galactose (for GST-dynein^MOTOR^), harvested, washed with cold water, and then resuspended in a small volume of water. The resuspended cell pellets were drop frozen into liquid nitrogen and then lysed in a coffee grinder. After lysis, 0.25 volume of 4X lysis buffer (1X buffer: 30 mM HEPES, pH 7.2, 50 mM potassium acetate, 2 mM magnesium acetate, 10% glycerol, 0.2 mM EGTA, 1 mM DTT, 0.1 mM Mg-ATP, 0.5 mM Pefabloc SC) was added, and the lysate was clarified at 22,000 x g for 20 min. The supernatant was then incubated with IgG sepharose 6 fast flow resin (GE) for 1 hour at 4°C, which was subsequently washed three times in lysis buffer, and twice in TEV buffer (50 mM Tris, pH 8.0, 150 mM potassium acetate, 2 mM magnesium acetate, 1 mM EGTA, 0.005% Triton X-100, 10% glycerol, 1 mM DTT, 0.1 mM Mg-ATP, 0.5 mM Pefabloc SC). The protein was then incubated in TEV buffer supplemented with TEV protease for 1 hour at 16°C. Following TEV digest, the solution was collected, and the resulting protein solution was aliquoted, flash frozen in liquid nitrogen, and stored at −80°C.

### Preparation of Zn^2+^/Chelator-buffered solutions

Buffers with precisely controlled Zn^2+^ concentrations were prepared according to previous established protocol^75^ for the *in vitro* motor ATPase assays. Solutions with Zn^2+^ from picomolar to micromolar concentrations were made by a pH-titration method. As indicated below, different Zn^2+^ chelators (EGTA and HEEDTA) and competing metal ions (Ca^2+^ or Sr^2+^) were used to buffer Zn^2+^ at pH 7.2 (for *in vitro* motility assays) or pH 6.9 (for *in vitro* microtubule polymerization assays).

To obtain free Zn^2+^ concentrations in the low picomolar range, we used HEEDTA with no competing metal ions. The stability constant of the Zn^2+^ complex of HEEDTA is: pK1 = 9.81, pK2 = 5.37, logK (ZnL) = 14.6. Two separate 100X stock solutions were made: 0.1 M HEEDTA (Buffer B4), and 0.1 M ZnCl_2_ + 0.1 M HEEDTA (Buffer A4). [HEEDTA]_t_ was set at 1 mM and [Zn^2+^]_t_ was varied from 0.1 – 0.9 mM in 0.1 mM increments by mixing 1X A4 and B4 buffers at different ratios to obtain the following free Zn^2+^ concentrations:

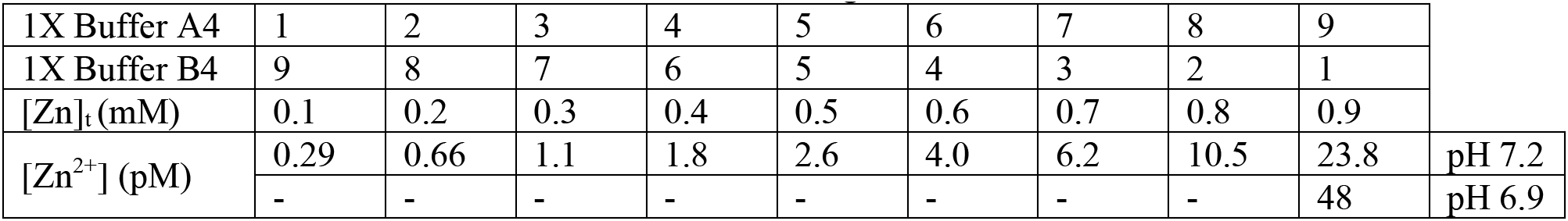

To obtain free Zn^2+^ concentrations in the high picomolar to low nanomolar range, we used EGTA with no competing metal ions. The stability constant of the Zn^2+^ complex of EGTA is pK1 = 9.40, pK2 = 8.78, logK(ZnL) = 12.6. Two separate 100X stock solutions were made: 0.1 M EGTA (Buffer B3), and 0.1 M ZnCl_2_ + 0.1 M EGTA (Buffer A3). [EGTA]_t_ was set at 1 mM and [Zn^2+^]_t_ was varied from 0.1 – 0.9 mM in 0.1 mM increments by mixing 1X A3 and B3 buffers at different ratios to obtain the following free Zn^2+^ concentrations:

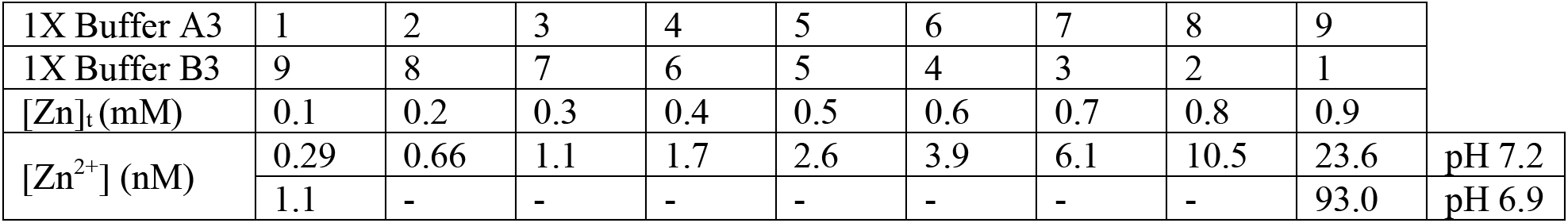

To obtain free Zn^2+^ concentrations in the nanomolar to low micromolar range, we used EGTA with Sr^2+^ as a competing metal ion. The stability constant of the Zn^2+^ complex of EGTA is: pK1 = 9.40, pK2 = 8.78, logK(SrL) = 8.50, logK(ZnL) = 12.6. Two separate 100X stock solutions were made: 0.1 M EGTA + 0.2 M SrCl_2_ (Buffer B2), and 0.1 M ZnCl_2_ + 0.1 M EGTA + 0.2 M SrCl_2_ (Buffer A2). [EGTA]_t_ and [Sr]_t_ was set at 1 mM, and [Zn^2+^]_t_ was varied from 0.1 – 0.9 mM in 0.1 mM increments by mixing 1X A2 and B2 buffers at different ratios to obtain the following free Zn^2+^ concentrations:

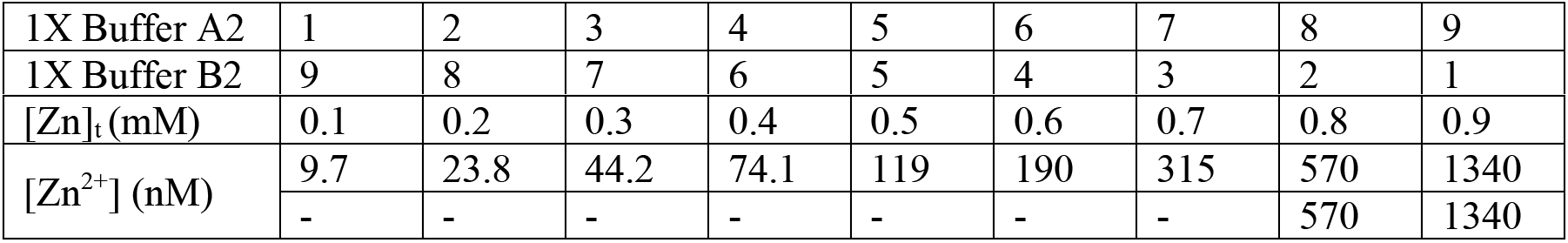

|  |  |  |  |  |  |  |  |  |  |
| --- | --- | --- | --- | --- | --- | --- | --- | --- | --- |
| 1X Buffer A2 | 1 | 2 | 3 | 4 | 5 | 6 | 7 | 8 | 9 |
| 1X Buffer B2 | 9 | 8 | 7 | 6 | 5 | 4 | 3 | 2 | 1 |

|  |  |  |  |  |  |  |  |  |  |  |
| --- | --- | --- | --- | --- | --- | --- | --- | --- | --- | --- |
| [Zn]<sub>t</sub> (mM) | 0.1 | 0.2 | 0.3 | 0.4 | 0.5 | 0.6 | 0.7 | 0.8 | 0.9 |  |
| [Zn<sup>2+</sup>] (nM) | 9.7 | 23.8 | 44.2 | 74.1 | 119 | 190 | 315 | 570 | 1340 | pH 7.2 |
|  | - | - | - | - | - | - | - | 570 | 1340 | pH 6.9 |

To obtain free Zn^2+^ concentrations in the micromolar range, we used EGTA with Ca^2+^ as a competing metal ion. The stability constant of Zn^2+^ complex of EGTA is: pK1 = 9.40, pK2 = 8.78, logK[CaL] = 10.86, logK(ZnL) = 12.6. Two separate 100X stock solutions were made: 0.1 M EGTA + 0.2 M CaCl_2_ (Buffer B1), and 0.1 M ZnCl_2_ + 0.1 M EGTA + 0.2 M CaCl_2_ (Buffer A1). [EGTA]_t_ and [Ca]_t_ was set at 1 mM, and [Zn^2+^]_t_ was varied from 0.1 – 0.9 mM in 0.1 mM increments by mixing 1X A1 and B1 buffers at different ratios to obtain the following free Zn^2+^ concentrations:

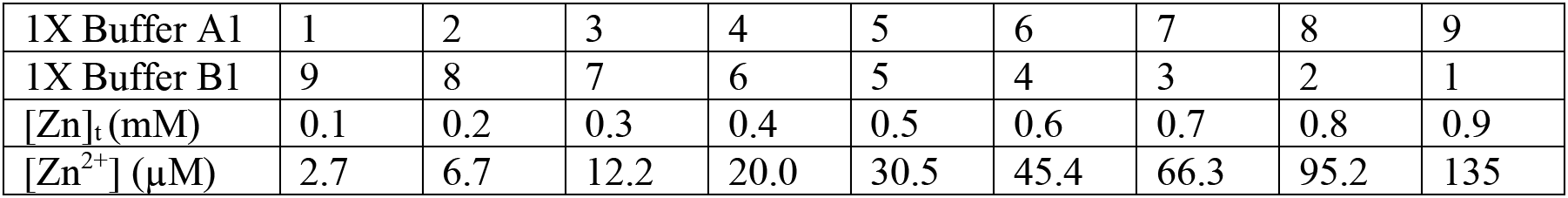

|  |  |  |  |  |  |  |  |  |  |
| --- | --- | --- | --- | --- | --- | --- | --- | --- | --- |
| 1X Buffer A1 | 1 | 2 | 3 | 4 | 5 | 6 | 7 | 8 | 9 |
| 1X Buffer B1 | 9 | 8 | 7 | 6 | 5 | 4 | 3 | 2 | 1 |
| [Zn]<sub>t</sub> (mM) | 0.1 | 0.2 | 0.3 | 0.4 | 0.5 | 0.6 | 0.7 | 0.8 | 0.9 |
| [Zn<sup>2+</sup>] (μM) | 2.7 | 6.7 | 12.2 | 20.0 | 30.5 | 45.4 | 66.3 | 95.2 | 135 |

### In vitro motor ATPase assays

For *in vitro* motor protein ATPase assays, the EnzChek^®^ Phosphate Assay Kit (#E-6646, Thermo Fisher Scientific, Waltham, MA) was used. Pre-formed, taxol-stabilized microtubules and recombinant human kinesin (KIF5A) were purchased from Cytoskeleton™ (Denver, CO). A minimally processive, artificially dimerized yeast dynein fragment (GST-dynein_331_) was purified as described above. Motor protein assay buffer (MPAB: 30 mM HEPES, pH 7.2, 50 mM potassium acetate, 2 mM magnesium acetate, 10% glycerol) was prepared using Chelex-treated H_2_O to remove all metal ion contaminants. 10 μM taxol, 500 μM DTT, 200 μM MESG, 1 U/mL purine nucleoside phosphorylase, 2 mM ATP, 1 μM taxol-stabilized microtubules, and 1x buffered zinc solutions (described above) were added to MPAB, then plated into a 96 well plate. Plates were spun down at 2250 rpm for 2 minutes, then incubated at room temperature for 10 minutes. Plates were read on a Synergy HTX Multi-Mode Microplate Reader (BioTek), collecting absorbance at 360 nm every 10 seconds. Background phosphate release was measured for 5 minutes, then either 5 nM GST-dynein_331_ or 10 nM KIF5A was added to each well, and absorbance at 360 nm was read for an additional 10 to 20 minutes. A phosphate standard was collected as described in the EnzChek Phosphate Assay Kit.

### In vitro microtubule polymerization assays

For *in vitro* microtubule polymerization assays, the traditional turbidity assay was used to monitor microtubule polymerization via absorbance at 340 nm with minor modifications. Purified porcine tubulin was purchased from Cytoskeleton™ (Denver, CO) and reconstituted to 10 mg/mL in microtubule assay buffer (“MTAB”: 80 mM PIPES, pH 6.9, 2 mM MgCl_2_, in Chelex-treated H_2_O) supplemented with 1 mM GTP. Microtubule reaction mix (MTAB, 5% glycerol, 2 mM GTP, and 5 mg/mL tubulin) was prepared on ice. 1X buffered Zn^2+^ solutions (described above) were plated into pre-chilled 96 well half-area plates, followed by microtubule reaction mix (50 μL total reaction volume). Plates were kept on ice until they were read on a Synergy HTX Multi-Mode Microplate Reader (BioTek), prewarmed to 37 °C, collecting absorbance at 340 nm every 20 seconds for 60 minutes.

### In vitro microtubule-Zn^2+^ binding assays

Pre-polymerized microtubules were purchased from Cytoskeleton™ (Denver, CO). Varying concentrations of microtubules were added to buffer (15 mM PIPES, 5 mM MgCl_2_, pH 7.0) supplemented with 10 μM taxol and 500 μM DTT. 1 μM ZnCl_2_ (prepared with Chelex-treated H_2_O) was then added, and the reaction was incubated for 10 minutes. 1 μM FluoZin-3 tetrapotassium salt (Thermo Fisher Scientific, Waltham, MA) was then added to each reaction, plated into a 96-well plate and incubated at room temperature for 10 minutes. Fluorescence was measured using a Synergy HTX Multi-Mode Microplate Reader (BioTek), collecting fluorescence every 30 seconds at 525 nm emission using a 488 nm excitation, for 5 minutes. As an internal control, 250 μM TPEN was added to each well to chelate all Zn^2+^ and plates were measures for 20 minutes, or until a fluorescent minimum plateau was reached. Data was normalized by subtracting the minimum post-TPEN fluorescence.

### In situ analysis of MAP interaction with microtubules

COS-7 cells were co-transfected with mCherry-α-tubulin and either EGFP-DCX, EGFP-Tau, EGFP-MAP2C, GFP-MAP1B, EGFP-p150Glued, mEmerald-MAP4, EGFP-MAP7, or EGFP-MAP9 and imaged 24 to 48 hours post-transfection on the Nikon/Solamere CSUX1 spinning disc confocal microscope as described above. Still images of several cells at recorded stage positions were taken before time-lapse imaging. Cells were imaged for a 5-minute baseline, then treated with metals and/or ionophores (as indicated) for 5 minutes. Still images of the same cells at the recorded stage positions were taken after the initial timelapse recording. For cells treated with 20 μM ZnCl_2_, Zn^2+^ was subsequently chelated with 100 μM TPEN and cells were imaged for 5 minutes. Finally, still images of the same cells at the recorded stage positions were taken after the TPEN treatment. Timelapse images were acquired every 10-20 seconds with a 50-200 ms exposure for both 488 nm and 561 nm laser excitation at 10 mW and 5 mW power, respectively.

The “Tubeness” analysis plugin for ImageJ was used to quantify microtubule morphology of MAPs, using a σ value of 1.5 pixels. Average tubeness for each cell was recorded at baseline, and each subsequent treatment, with values normalized to the baseline tubeness for each cell.

For depolarization experiments, primary hippocampal neurons (DIV 10 to 16) were depolarized for 5 minutes with 50 mM KCl in the presence or absence of 100 μM ZnCl_2_ in HHBSS lacking Ca^2+^. Neurons were immediately fixed with ice-cold methanol for 10 minutes at −20 °C to simultaneously permeabilize neurons and fix cells in a manner which would preserve microtubules and MAP interactions on microtubules. Fixed cells were blocked with 3% BSA for 30 minutes, then immediately immunostained (no PBS wash) with anti-Tau polyclonal primary antibody (reactive to human, mouse, rat) raised in rabbit (IgG isotype) at a 1:700 dilution for 1 hour at room temperature or overnight at 4 °C. Cells were then washed with PBS, and re-blocked with 3% BSA for 15 minutes. Cells were then incubated with goat anti-Rabbit IgG (H+L) Cross-Adsorbed secondary antibody conjugated to Alexa Fluor™ 488 (A-11008) at a 1:500 dilution and beta Tubulin loading control monoclonal antibody conjugated to Alexa Fluor™ 555 (MA5-16308) at a 1:250 dilution for 45 minutes. Cells were finally washed with PBS then stained with 600 nM DAPI and imaged on the Nikon/Solamere CSUX1 spinning disc confocal microscope as described above. Multichannel images were taken across a 1 mm^2^ area of the neuron culture for each dish and stitched together using the built-in Fiji “Stiching” plugin. Custom code (ImageJ Macro Language) was used to analyze the individual images to segment microtubule-positive areas, primarily axons, and measure average fluorescence intensity of Alexa Fluor™ 488.

### Analysis of MAP interaction and predicted Zn^2+^ binding sites on microtubules

Molecular graphics and analyses were performed with UCSF Chimera^76^, developed by the Resource for Biocomputing, Visualization, and Informatics at the University of California, San Francisco, with support from NIH P41-GM103311.

Previously published cryo-EM microtubule reconstruction models (PDB ID: 6EVY) were used for MAP interaction and Zn^2+^ binding prediction analysis. The only available cryo-EM reconstruction models of microtubule-bound MAPs analyzed were DCX (PDB ID: 6REV), Tau (PDB ID: 6CVN), kinesin microtubule binding domain (PDB ID: 3J8X), and cytoplasmic dynein microtubule binding domain (PDB ID: 6RZB). Individually, the microtubule-bound MAPs were analyzed using the Protein-Ligand Interaction Profiler (PLIP)^42^, where the MAP was specified as the ligand and all inter-chain interactions (hydrophobic interactions, hydrogen bonds, and salt bridges) between the ligand and α-tubulin or β-tubulin were analyzed. Using Chimera, α-tubulin and β-tubulin residues identified by PLIP were colored to correspond with the MAP(s) that interacted with the residue.

Zn^2+^ binding prediction analysis of α-tubulin or β-tubulin was performed using the MIB (Metal Ion-Binding Site Prediction and Docking) Server^44^ Chain A (α-tubulin) and Chain B (β-tubulin) from the cryo-EM microtubule reconstruction model (PDB ID: 6EVY) were run individually. α-tubulin and β-tubulin residues identified by MIB with a score > 0 were colored using the “Render by Attribute” interface.

### Data analysis and statistics

Imaging data were analyzed with Fiji (ImageJ, National Institutes of Health) and raw data output from Fiji were analyzed using Microsoft Excel in combination with JMP software (JMP^®^, Version 15.2.0. SAS Institute Inc., Cary, NC), and Mathematica (Wolfram Research, Inc., Version 12.1, Champaign, IL (2020)). Statistical analysis was performed using Excel or JMP software, using the appropriate statistical tests. All unpaired t-tests were preceded by an F-test to analyze variance equality. t-tests were then performed with consideration for the determined variance equality or inequality. Least-squares regression with comparisons across all groups used restricted maximum likelihood estimation (REML) and post-hoc multiple comparisons used Tukey HSD. All measurements were taken from distinct samples. No region of interest (ROI) was measured repeatedly. When selecting a ROI in the soma for cytosolic sensors that were also present in the nucleus, care was taken to avoid selecting nuclear areas. For time traces of fluorescence intensity, background fluorescence was subtracted from ROIs. For single wavelength sensors, changes in fluorescence intensity (ΔF = F-F_0_) were normalized to the baseline preceding the 0 second time point (F0), indicated as ΔF/F_0_.

## Supplemental Information

### Supplemental Tables

**Supplementary Table 1:** Statistical analysis of ion effects on DCX decoration of microtubules. Listed below are the *p*-values and associated connecting letters report from the post-hoc Tukey HSD test between various ion influx treatments of COS-7 cells expressing EGFP-DCX. Bolded values indicate statistically significant *p*-values (below α = 0.05). Related to Figure 5E.

| Comparison | *p* | Connecting Letters Report |  |  |
| --- | --- | --- | --- | --- |
|  |  | Group | Letters | Mean |
| 2 mM CaCl<sub>2</sub> vs. 20 μM ZnCl<sub>2</sub> | **<.0001** | 2 mM CaCl<sub>2</sub> | A | 1.098973 |
| 20 μM CuCl<sub>2</sub> vs. 20 μM ZnCl<sub>2</sub> | **<.0001** | 20 μM CuCl<sub>2</sub> | A | 1.076604 |
| 100 μM CuCl<sub>2</sub> vs. 20 μM ZnCl<sub>2</sub> | **<.0001** | 100 μM CuCl<sub>2</sub> | A | 1.026417 |
| 2.5 μM PTO vs. 20 μM ZnCl<sub>2</sub> | **<.0001** | 2.5 μM PTO | A | 0.995647 |
| 2 mM CaCl<sub>2</sub> vs. 2.5 μM PTO | 0.6877 | 20 μM ZnCl<sub>2</sub> | B | 0.596121 |
| 20 μM CuCl<sub>2</sub> vs. 2.5 μM PTO | 0.8436 | | | |
| 2 mM CaCl<sub>2</sub> vs. 100 μM CuCl<sub>2</sub> | 0.8789 | | | |
| 20 μM CuCl<sub>2</sub> vs. 100 μM CuCl<sub>2</sub> | 0.9657 | | | |
| 100 μM CuCl<sub>2</sub> vs. 2.5 μM PTO | 0.9948 | | | |
| 2 mM CaCl<sub>2</sub> vs. 20 μM CuCl<sub>2</sub> | 0.9985 | | | |

**Supplementary Table 2:**
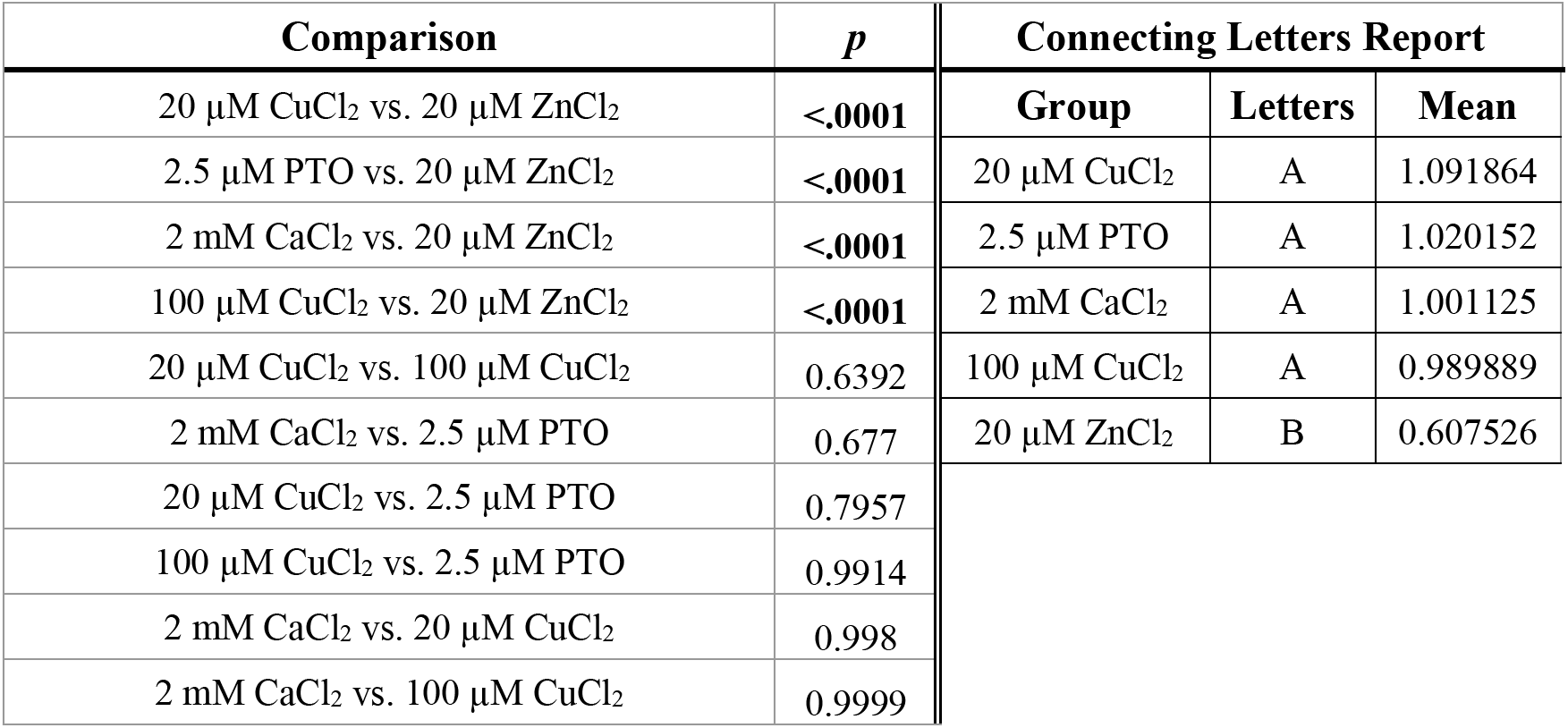
Statistical analysis of ion effects on Tau decoration of microtubules. Listed below are the *p*-values and associated connecting letters report from the post-hoc Tukey HSD test between various ion influx treatments of COS-7 cells expressing EGFP-Tau. Bolded values indicate statistically significant *p*-values (below α = 0.05). Related to Figure 5F.

**Supplementary Table 3:** Statistical analysis of ion effects on MAP2C decoration of microtubules. Listed below are the *p*-values and associated connecting letters report from the post-hoc Tukey HSD test between various ion influx treatments of COS-7 cells expressing EGFP-MAP2C. Bolded values indicate statistically significant *p*-values (below α = 0.05). Related to Figure 5G.

| Comparison | $p$ | Connecting Letters Report | | |
| --- | --- | --- | --- | --- |
|  |  | Group | Letters | Mean |
| 2.5 μM PTO vs. 20 μM ZnCl<sub>2</sub> | **<.0001** | 2.5 μM PTO | A | 1.139288 |
| 20 μM CuCl<sub>2</sub> vs. 20 μM ZnCl<sub>2</sub> | **0.0003** | 20 μM CuCl<sub>2</sub> | A | 1.025192 |
| 2 mM CaCl<sub>2</sub> vs. 20 μM ZnCl<sub>2</sub> | **0.0017** | 2 mM CaCl<sub>2</sub> | A | 1.009978 |
| 100 μM CuCl<sub>2</sub> vs. 20 μM ZnCl<sub>2</sub> | **0.0026** | 100 μM CuCl<sub>2</sub> | A | 0.994818 |
| 100 μM CuCl<sub>2</sub> vs. 2.5 μM PTO | 0.0556 | 20 μM ZnCl<sub>2</sub> | B | 0.805633 |
| 2 mM CaCl<sub>2</sub> vs. 2.5 μM PTO | 0.1365 | | | |
| 20 μM CuCl<sub>2</sub> vs. 2.5 μM PTO | 0.2036 | | | |
| 20 μM CuCl<sub>2</sub> vs. 100 μM CuCl<sub>2</sub> | 0.9794 | | | |
| 2 mM CaCl<sub>2</sub> vs. 20 μM CuCl<sub>2</sub> | 0.9987 | | | |
| 2 mM CaCl<sub>2</sub> vs. 100 μM CuCl<sub>2</sub> | 0.9988 | | | |

**Supplementary Table 4:**
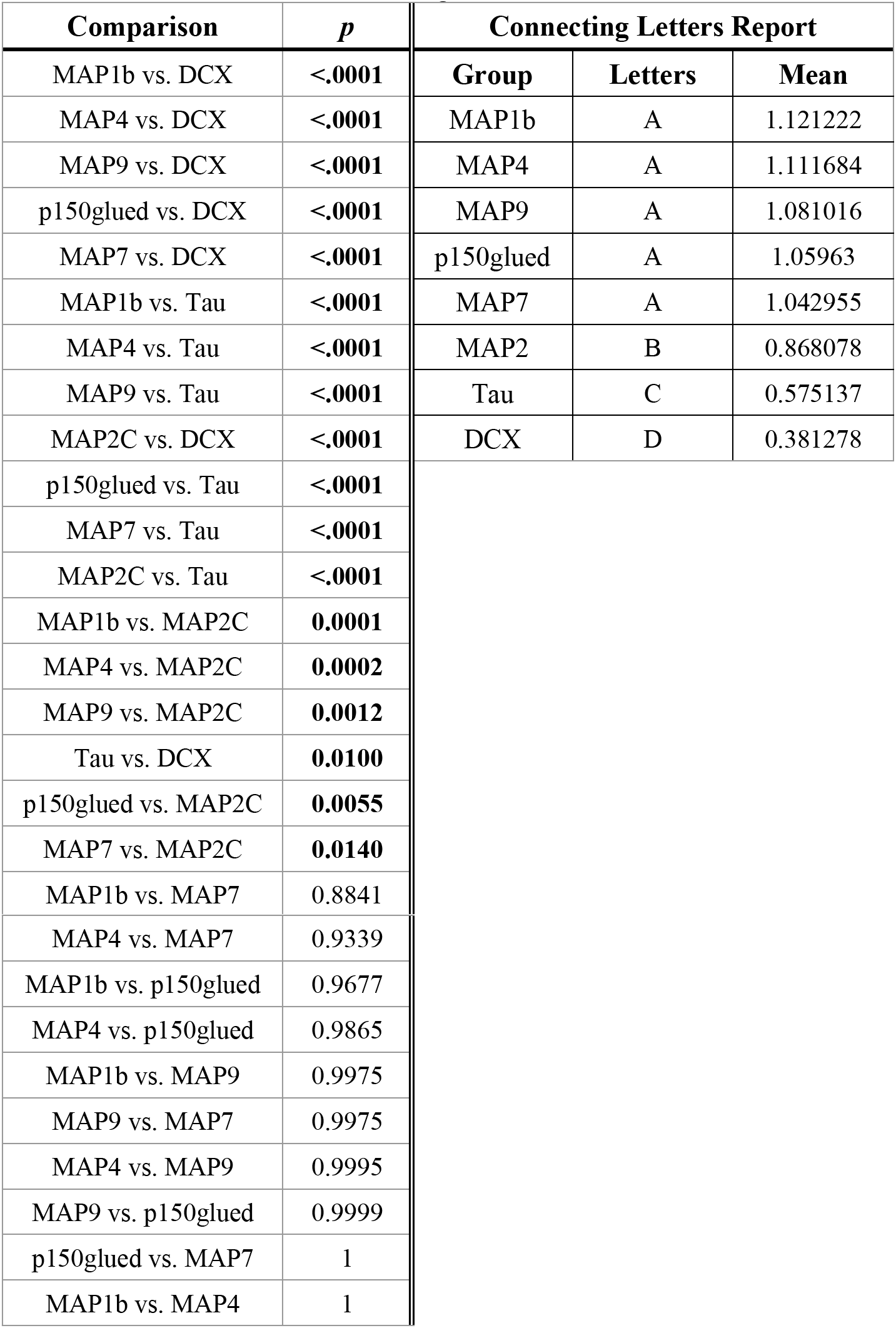
Statistical analysis of Zn^2+^ effect on MAP decoration of microtubules. Listed below are the *p*-values and associated connecting letters report from the post-hoc Tukey HSD test between MAP1b, MAP2C, MAP4, MAP7, MAP9, p150glued, DCX, and Tau. Bolded values indicate statistically significant *p*-values (below α = 0.05). Related to Figure 5H.

| Comparison | $p$ | Connecting Letters Report | | |
| --- | --- | --- | --- | --- |
|  |  | Group | Letters | Mean |
| MAP1b vs. DCX | **<.0001** | MAP1b | A | 1.121222 |
| MAP4 vs. DCX | **<.0001** | MAP4 | A | 1.111684 |
| MAP9 vs. DCX | **<.0001** | MAP9 | A | 1.081016 |
| p150glued vs. DCX | **<.0001** | p150glued | A | 1.05963 |
| MAP7 vs. DCX | **<.0001** | MAP7 | A | 1.042955 |
| MAP1b vs. Tau | **<.0001** | MAP2 | B | 0.868078 |
| MAP4 vs. Tau | **<.0001** | Tau | C | 0.575137 |
| MAP9 vs. Tau | **<.0001** | DCX | D | 0.381278 |
| MAP2C vs. DCX | **<.0001** |  |  |  |
| p150glued vs. Tau | **<.0001** |  |  |  |
| MAP7 vs. Tau | **<.0001** |  |  |  |
| MAP2C vs. Tau | **<.0001** |  |  |  |
| MAP1b vs. MAP2C | **0.0001** |  |  |  |
| MAP4 vs. MAP2C | **0.0002** |  |  |  |
| MAP9 vs. MAP2C | **0.0012** |  |  |  |
| Tau vs. DCX | **0.0100** |  |  |  |
| p150glued vs. MAP2C | **0.0055** |  |  |  |
| MAP7 vs. MAP2C | **0.0140** |  |  |  |
| MAP1b vs. MAP7 | 0.8841 |  |  |  |

### Supplemental Figures

**Figure S1:**
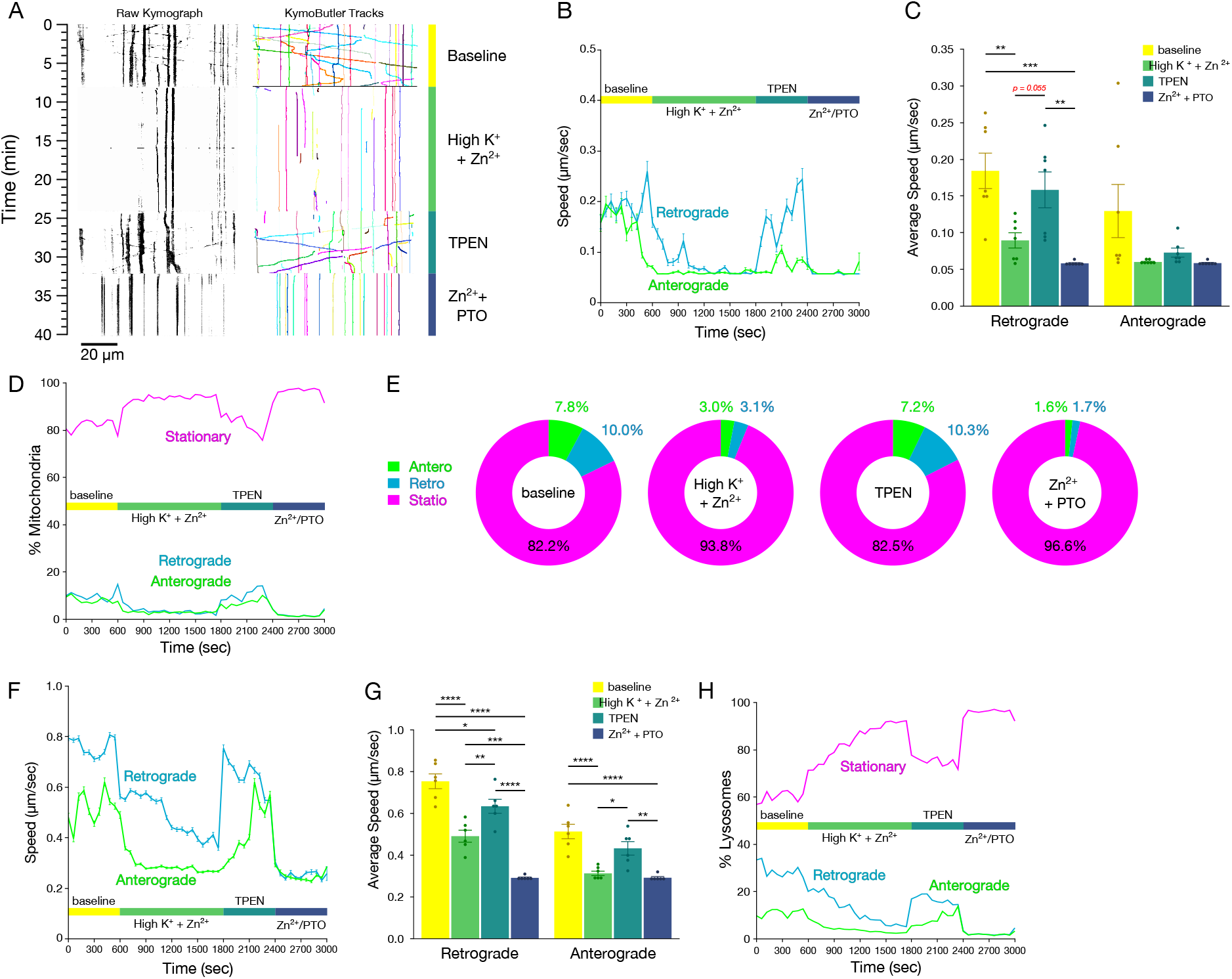
Zn^2+^ influx arrests axonal transport of mitochondria and lysosomes. (A) Representative kymographs (left) and measured tracks (identified using KymoButler; right) of mito-mCherry motility along axons from primary cultured rat hippocampal neurons prior to (“baseline”; top) or after initiation of Zn^2+^ influx by high K^+^ depolarization (middle-top), following washout and addition of 100 μM TPEN (middle-bottom), and then washout and addition of 100 μM Zn^2+^ and 2.5 μM PTO (bottom). (B) Mean instantaneous speed (±S.E.M.) of mitochondria moving retrograde (blue) or anterograde (green) across baseline, Zn^2+^ influx by high K^+^ depolarization, TPEN, and Zn^2+^ + PTO treatment (60-second binned, representing 7 axons). (C) Mean speed (±S.E.M.) of mitochondria moving in the indicated directions for each condition (n = 7 axons from 3 independent replicates). (D) Proportions of mitochondria moving in the indicated directions across baseline, depolarization, TPEN, and Zn^2+^ + PTO treatment (60-second binned, representing 7 axons). (E) Mean proportions of mitochondrial motility for each condition. (F) Mean instantaneous speed (±S.E.M.) of lysosomes moving retrograde (blue) or anterograde (green) across baseline, Zn^2+^ influx by high K^+^ depolarization, following washout and addition of 100 μM TPEN, and then washout and addition of 100 μM Zn^2+^ and 2.5 μM PTO (bottom). (60-second binned, representing 6 axons). (G) Mean speed (±S.E.M.) of lysosomes moving in the indicated directions for each condition (n = 6 axons from 3 independent replicates). (H) Proportions of lysosomes moving in the indicated directions across baseline, Zn^2+^ influx by high K^+^ depolarization, TPEN, and Zn^2+^ + PTO treatment (60-second binned, representing 6 axons). All experiments were performed in the absence of extracellular Ca^2+^. **** *p* < 0.0001, *** *p* < 0.001, ** *p* < 0.01, * *p* < 0.05, n.s. not significant.

**Figure S2:**
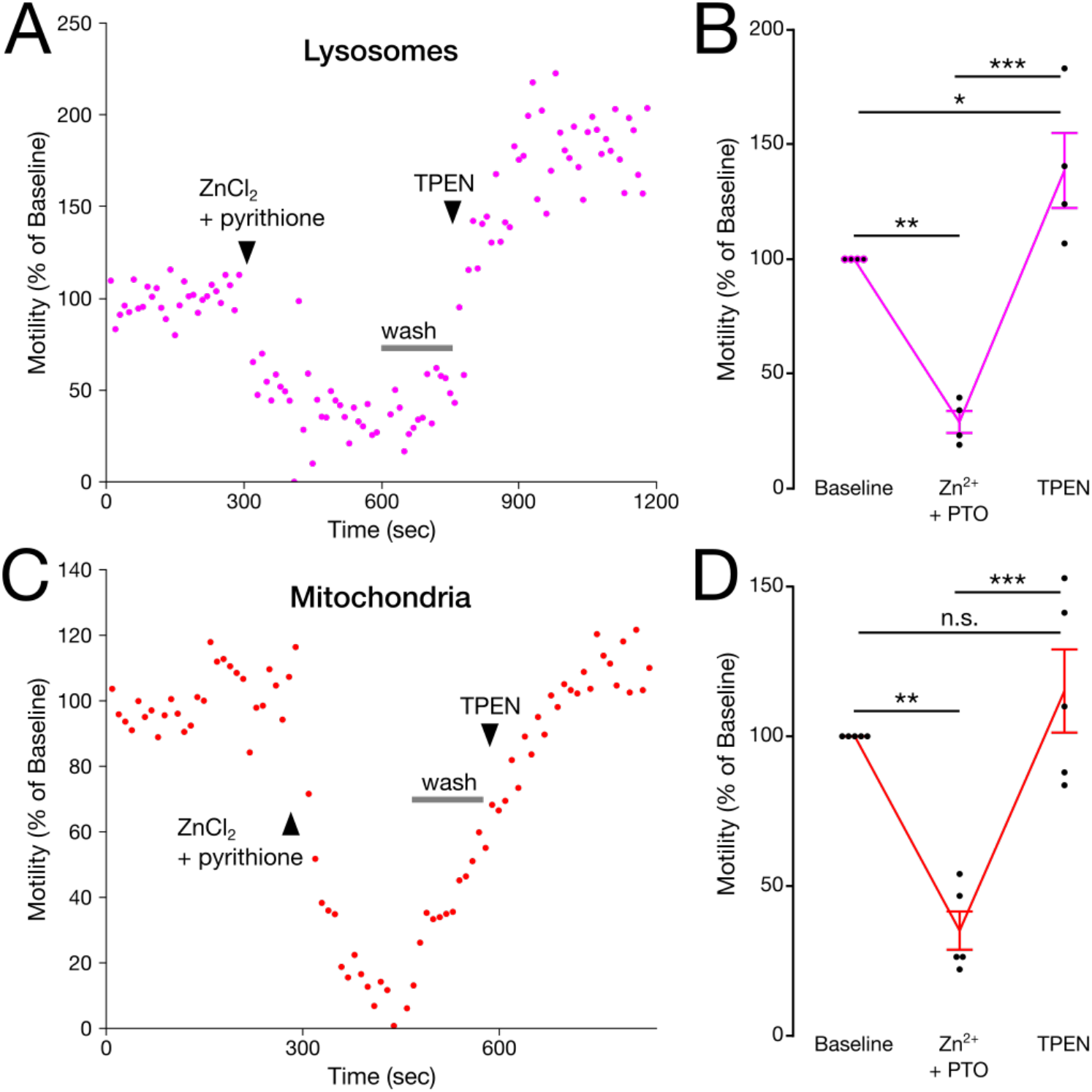
Zn^2+^ influx inhibits motility of lysosomes and mitochondria in HeLa cells. (A and C) Representative lysosomal (A) and mitochondrial (C) motility (determined using the Total Motility plugin for Fiji; see Methods) in HeLa cells treated with 20 μM ZnCl_2_ and 1.25 μM pyrithione (PTO), then washed and treated with 100 μM TPEN (treatments added at time points indicated on plots). (B and D) Mean (±S.E.M.) lysosomal (B) and mitochondrial (D) motility normalized to their respective baselines (n = 4 and 5 cells, from 4 and 3 independent replicates, respectively). One-way repeated measures ANOVA, with post-hoc Tukey HSD. All experiments were performed in the absence of extracellular Ca^2+^. *** *p* < 0.0001, ** *p* < 0.01, * *p* < 0.05, n.s. not significant.

**Figure S3:**
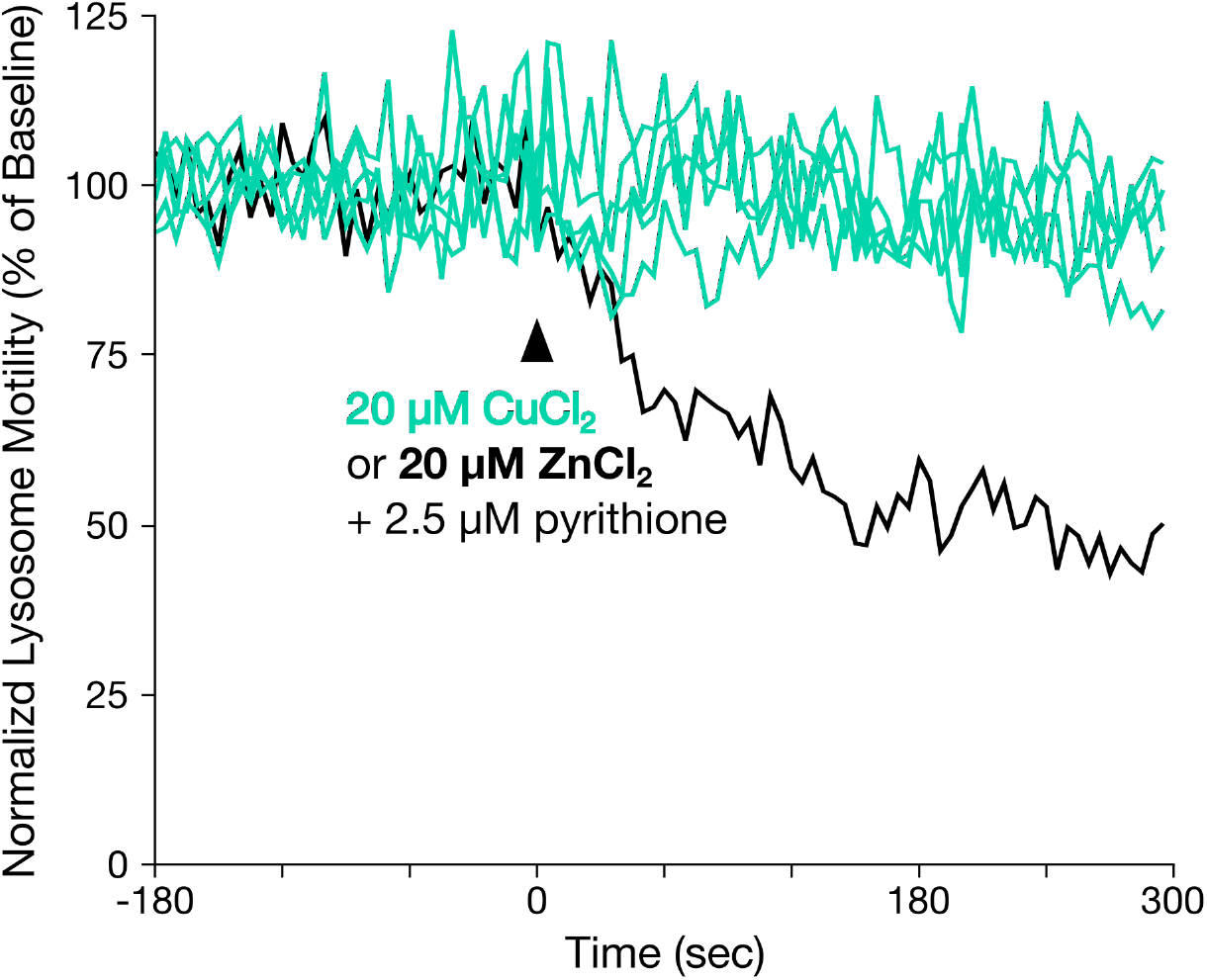
Cu^2+^ has no effect on lysosomal motility in HeLa cells. (A) Representative traces of lysosomal motility (determined using the Total Motility plugin for Fiji; see Methods) in HeLa cells stained with LysoTracker Red and treated with 2.5 μM pyrithione and either 20 μM CuCl_2_ (teal lines) or 20 μM ZnCl_2_ (black line) at 0 sec. All experiments were performed in the absence of extracellular Ca^24^.

**Figure S4:**
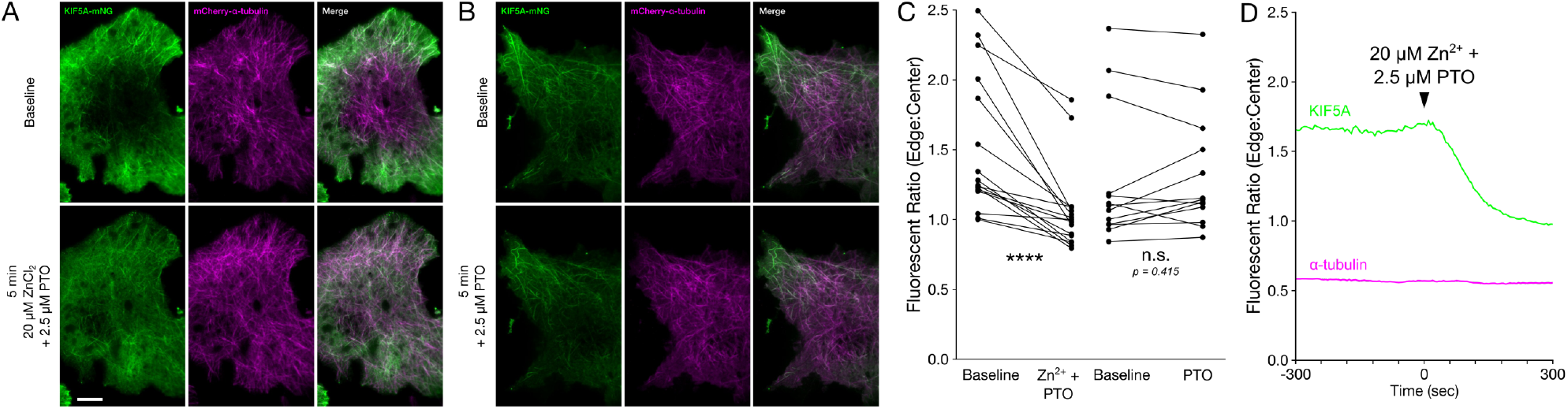
Zn^2+^ can bind microtubules and inhibits motor protein activity in situ and in vitro. (A-B) Representative oblique illumination micrographs of COS-7 cells expressing KIF5A(1-560)-mNG-FRB (green, left) and mCherry-α-tubulin (magenta, middle), and merged channels (right). Cells were imaged at baseline (top) and 5 minutes after treatment (bottom) with (A) 20 μM ZnCl_2_ and 2.5 μM pyrithione (PTO), or (B) 2.5 μM PTO alone. (C) Ratio of edge-to-center fluorescence of KIF5A(1-560)-mNG-FRB in COS-7 cells as treated in (A) and (B) (Zn^2+^-treated, n = 16 cells from 3 individual biological replicates; PTO control, n = 13 cells from 3 individual biological replicates). Two-tailed paired t-tests. (D) Representative timelapse of edge-to-center fluorescence of KIF5A(1-560)-mNG-FRB (green) and mCherry-α-tubulin (magenta) in COS-7 cells, treated as in (A). All *in situ* experiments were performed in the absence of extracellular Ca^2+^. \*\*\*\**p* < 0.0001, n.s. not significant.

**Figure S5:**
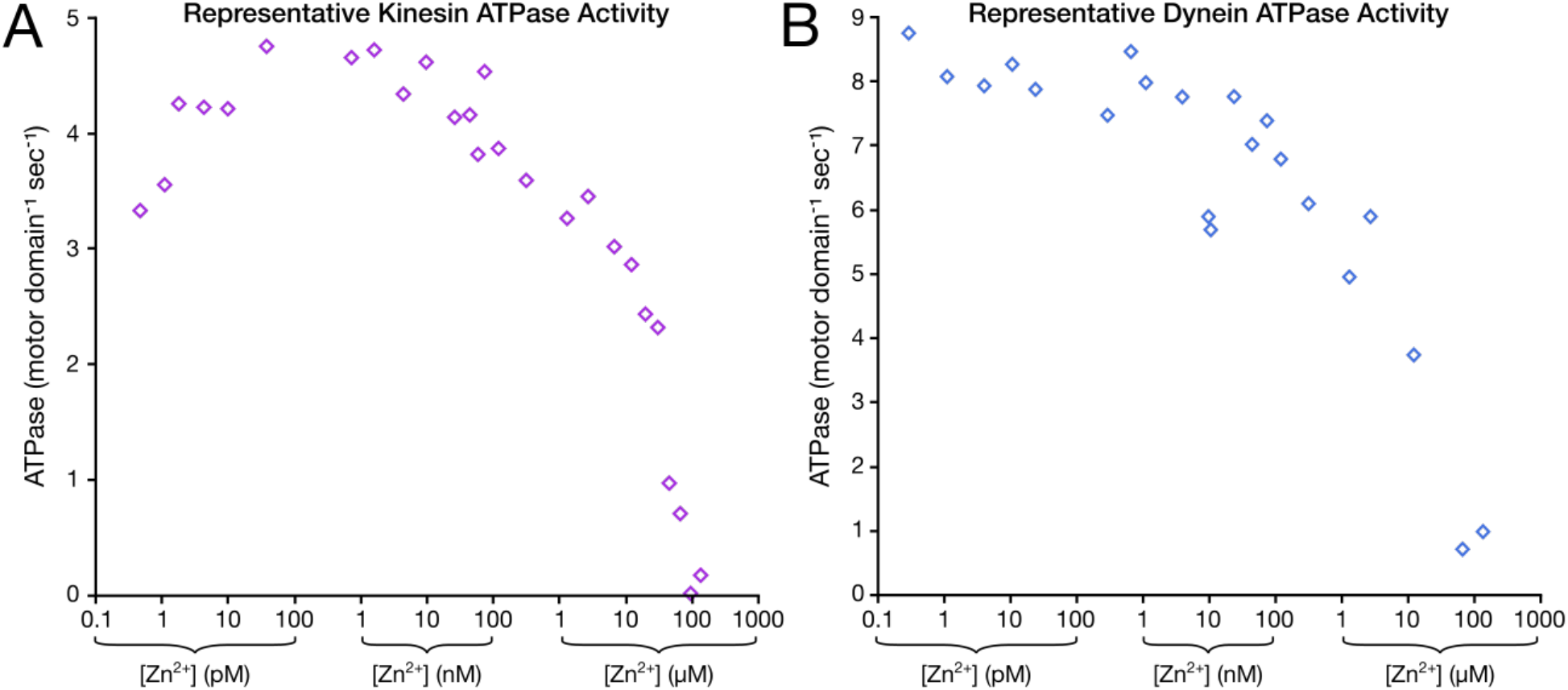
Supplemental in vitro analysis of the effect of Zn^2+^ on kinesin and dynein motor activity. (A-B) Representative microtubule-stimulated ATPase activity showing non-normalized ATPase activity per motor domain per second for purified (A) recombinant human kinesin (KIF5A) or (B) a minimally processive, artificially dimerized yeast dynein fragment (GST-dynein_331_) across a range of Zn^2+^ concentrations (from same dataset as Figure 4A-B).

**Figure S6:**
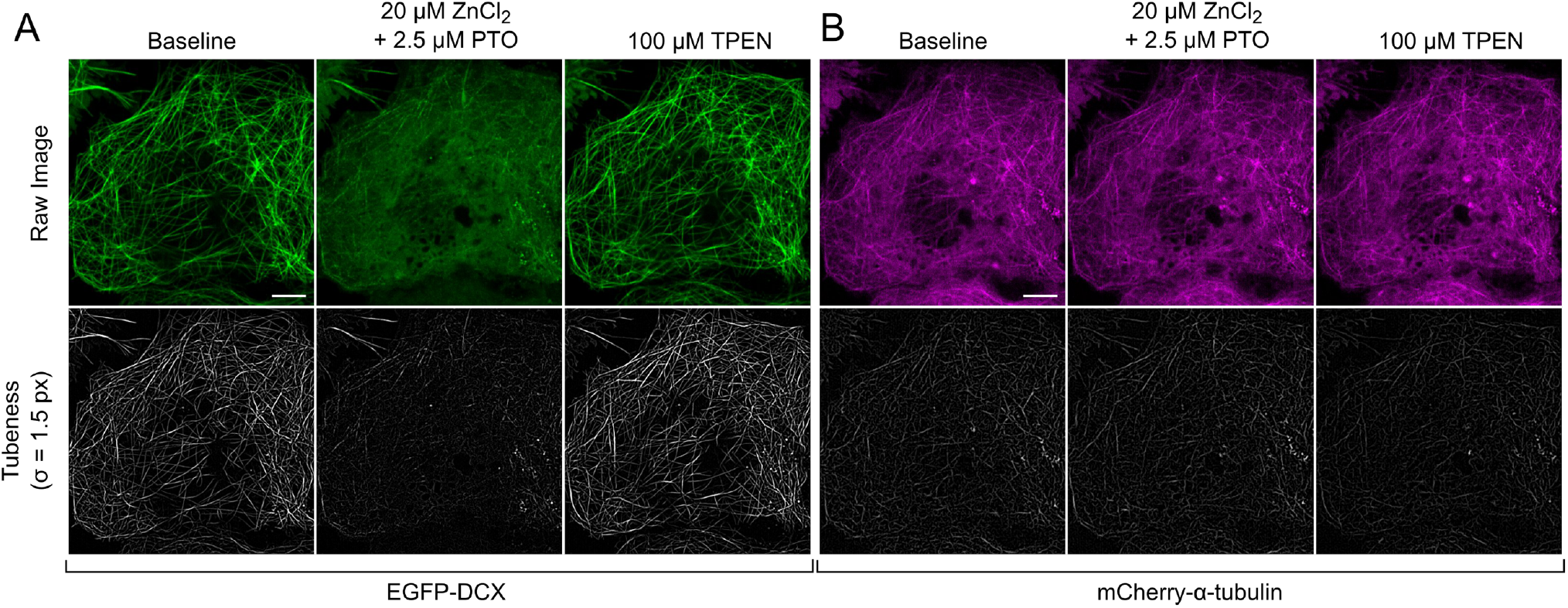
Tubeness analysis of EGFP-DCX expressing COS-7 cells with Zn^2+^ and TPEN treatment. (A-B) Representative micrographs of COS-7 cells expressing (A) EGFP-DCX and (B) mCherry-α-tubulin before treatment (“baseline”, left), after 5-minute treatment with 20 μM ZnCl_2_ and 2.5 μM PTO (center), or after subsequent treatment with 100 μM TPEN (right). Scale bar = 10 μm. Raw fluorescence images (top) and “Tubeness” analysis (Fiji Plugin) with a sigma, σ, of 1.5 pixels (bottom) are shown.

**Figure S7:**
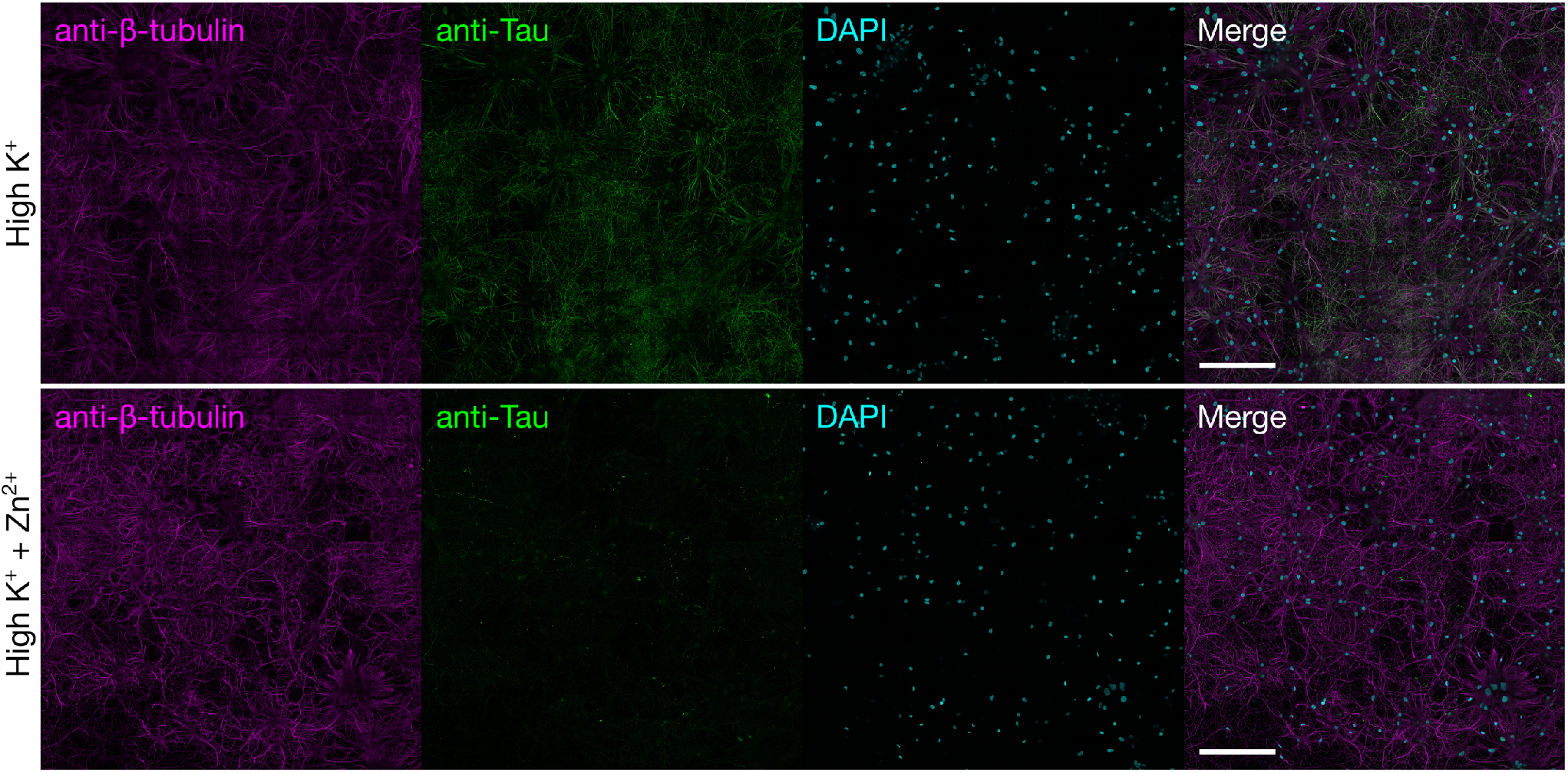
Depolarization induced Zn^2+^ influx promotes detachment of endogenous neuronal Tau in situ. Representative immunofluorescence micrographs (15 x 15 stitched image grid, 5% overlap) of methanol-fixed primary rat hippocampal neurons depolarized with 50 mM KCl in the absence (top) or presence (bottom) of 100 μM ZnCl_2_. Fixed cells were incubated with anti-β-tubulin monoclonal antibodies (magenta, left), anti-Tau polyclonal antibodies (green, left center), DAPI (cyan, right center), and channels are shown merged (right). Scale bar = 200 μm. All experiments were performed in the absence of extracellular Ca^2+^.

**Figure S8:**
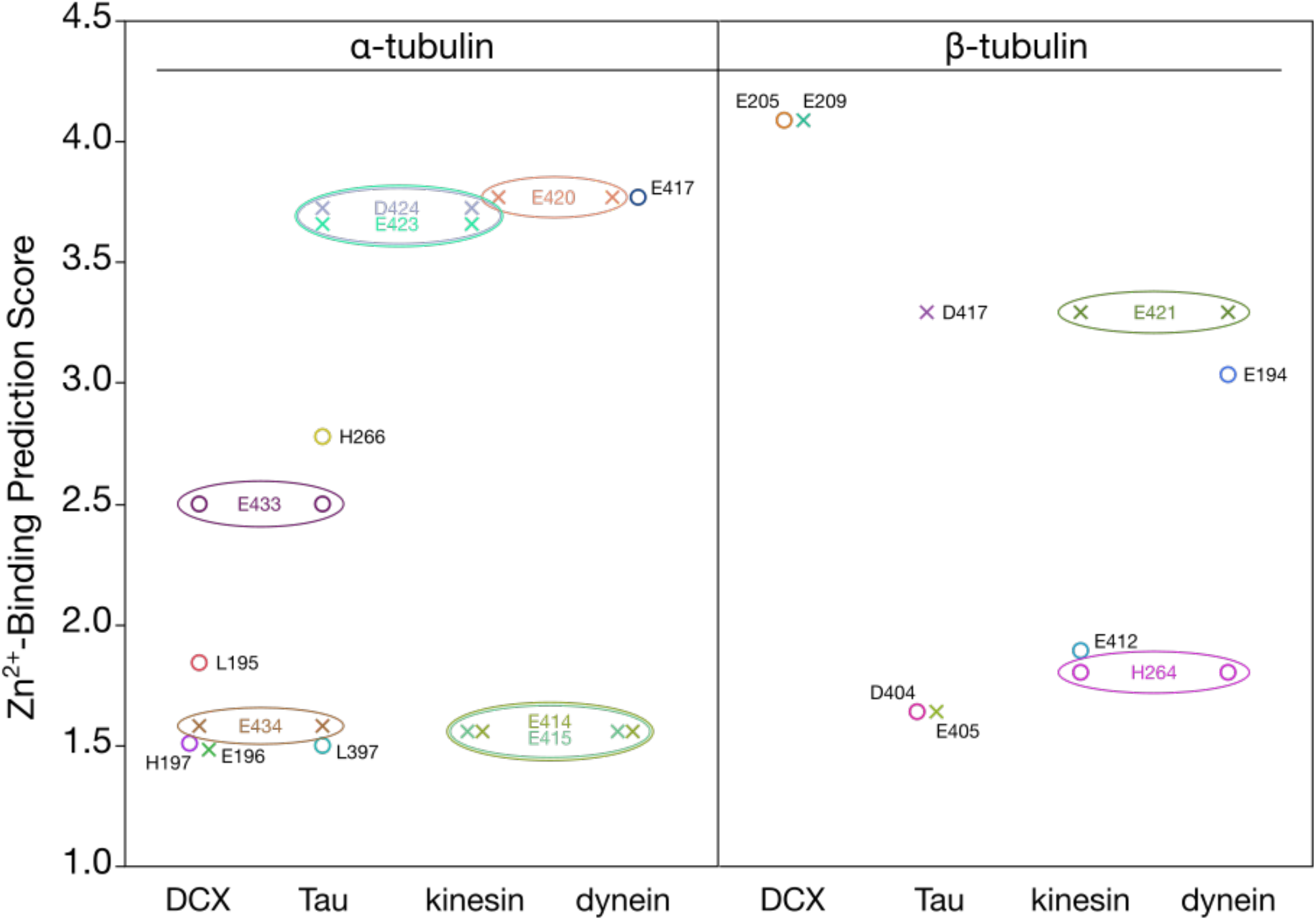
Predicted Zn^2+^ binding near MAP binding sites. Zn^2+^-binding prediction scores of α-tubulin and β-tubulin amino acid residues that directly interact with the indicated MAPs (“X” markers) and residues within 3Å of MAP-interacting residues (open circle markers). Residues that are shared between MAPs are represented by enclosed ovals.

### Supplemental Video Captions

**Video S1: *Depolarization-induced influx of Zn^2+^ arrests axonal transport of lysosomes.*** LAMP1-mCherry labeled lysosomes moving along axons of primary rat hippocampal neurons during baseline, depolarization induced Zn^2+^ influx (High K + Zn^2+^), and washout followed by TPEN treatment. Related to Figure 1.

**Video S2: *Depolarization-induced influx of Ca^2+^does not inhibit axonal transport of lysosomes.*** LAMP1-mCherry labeled lysosomes moving along axons of primary rat hippocampal neurons during baseline, depolarization induced Ca^2+^ influx (High K + Ca^2+^), and washout. Related to Figure 1.

**Video S3: *Zn^2+^ influx arrests axonal transport of mitochondria.*** mito-mCherry labeled mitochondria moving along axons of primary rat hippocampal neurons during baseline, depolarization induced Zn^2+^ influx (High K + Zn^2+^), washout followed by TPEN treatment, and washout followed by Zn^2+^ and pyrithione (PTO) treatment. Related to Figure S1.

**Video S4: *Zn^2+^ inhibits organellar motility with nanomolar IC_50_ in HeLa cells.*** Simultaneous imaging of Zn^2+^ (using GZnP2 sensor) and either lysosome (LAMP1-mCherry) or mitochondria (mito-mCherry) motility in HeLa cells. Related to Figure 2 and Figure S2.

**Video S5: *Zn^2+^ inhibits KIF5A movement in situ in a dose dependent manner***. Peroxisome dispersion assays in COS-7 cells expressing KIF5A(1-560)-mNG-FRB and PEX-mRFP-2xFKBP, during baseline, treatment with pyrithione (PTO) and varying concentrations of Zn^2+^, and subsequent treatment with Zotarolimus. Related to Figure 3.

**Video S6: *Zn^2+^ redistributes KIF5A in COS-7 cells.*** Distribution of KIF5A(1-560)-mNG-FRB and mCherry-α-tubulin in COS-7 cells during baseline and after Zn^2+^ and pyrithione (PTO) treatment. Related to Figure S4.

**Video S7: *Zn^2+^ promotes the dissociation of doublecortin from microtubules in situ.*** Microtubule decoration of EGFP-DCX in COS-7 cells during baseline, Zn^2+^ and pyrithione (PTO) treatment, and Zn^2+^ chelation with 100 μM TPEN. Related to Figure 5.

