## Supplemental information for "Zinc arrests axonal transport and displaces tau, doublecortin, and MAP2C from microtubules"

**Supplemental Tables**

**Supplementary Table 1: *Statistical analysis of ion effects on DCX decoration of microtubules.*** Listed below are the *p*-values and associated connecting letters report from the post-hoc Tukey HSD test between various ion influx treatments of COS-7 cells expressing EGFP-DCX. Bolded values indicate statistically significant *p*-values (below α = 0.05). Related to Figure 5E.

| Comparison | *p* | Connecting Letters Report | | |
| --- | --- | --- | --- | --- |
| 2 mM CaCl_2_ vs. 20 µM ZnCl_2_ | **<.0001** | **Group** | **Letters** | **Mean** |
| 20 µM CuCl_2_ vs. 20 µM ZnCl_2_ | **<.0001** | 2 mM CaCl_2_ | A | 1.098973 |
| 100 µM CuCl_2_ vs. 20 µM ZnCl_2_ | **<.0001** | 20 µM CuCl_2_ | A | 1.076604 |
| 2.5 µM PTO vs. 20 µM ZnCl_2_ | **<.0001** | 100 µM CuCl_2_ | A | 1.026417 |
| 2 mM CaCl_2_ vs. 2.5 µM PTO | 0.6877 | 2.5 µM PTO | A | 0.995647 |
| 20 µM CuCl_2_ vs. 2.5 µM PTO | 0.8436 | 20 µM ZnCl_2_ | B | 0.596121 |
| 2 mM CaCl_2_ vs. 100 µM CuCl_2_ | 0.8789 |  |  |  |
| 20 µM CuCl_2_ vs. 100 µM CuCl_2_ | 0.9657 |  |  |  |
| 100 µM CuCl_2_ vs. 2.5 µM PTO | 0.9948 |  |  |  |
| 2 mM CaCl_2_ vs. 20 µM CuCl_2_ | 0.9985 |  |  |  |

**Supplementary Table 2: *Statistical analysis of ion effects on Tau decoration of microtubules.*** Listed below are the *p*-values and associated connecting letters report from the post-hoc Tukey HSD test between various ion influx treatments of COS-7 cells expressing EGFP-Tau. Bolded values indicate statistically significant *p*-values (below α = 0.05). Related to Figure 5F.

| Comparison | *p* | Connecting Letters Report | | |
| --- | --- | --- | --- | --- |
| 20 µM CuCl_2_ vs. 20 µM ZnCl_2_ | **<.0001** | **Group** | **Letters** | **Mean** |
| 2.5 µM PTO vs. 20 µM ZnCl_2_ | **<.0001** | 20 µM CuCl_2_ | A | 1.091864 |
| 2 mM CaCl_2_ vs. 20 µM ZnCl_2_ | **<.0001** | 2.5 µM PTO | A | 1.020152 |
| 100 µM CuCl_2_ vs. 20 µM ZnCl_2_ | **<.0001** | 2 mM CaCl_2_ | A | 1.001125 |
| 20 µM CuCl_2_ vs. 100 µM CuCl_2_ | 0.6392 | 100 µM CuCl_2_ | A | 0.989889 |
| 2 mM CaCl_2_ vs. 2.5 µM PTO | 0.677 | 20 µM ZnCl_2_ | B | 0.607526 |
| 20 µM CuCl_2_ vs. 2.5 µM PTO | 0.7957 |  |  |  |
| 100 µM CuCl_2_ vs. 2.5 µM PTO | 0.9914 |  |  |  |
| 2 mM CaCl_2_ vs. 20 µM CuCl_2_ | 0.998 |  |  |  |
| 2 mM CaCl_2_ vs. 100 µM CuCl_2_ | 0.9999 |  |  |  |

**Supplementary Table 3: *Statistical analysis of ion effects on MAP2C decoration of microtubules.*** Listed below are the *p*-values and associated connecting letters report from the post-hoc Tukey HSD test between various ion influx treatments of COS-7 cells expressing EGFP-MAP2C. Bolded values indicate statistically significant *p*-values (below α = 0.05). Related to Figure 5G.

| Comparison | *p* | Connecting Letters Report | | |
| --- | --- | --- | --- | --- |
| 2.5 µM PTO vs. 20 µM ZnCl_2_ | **<.0001** | **Group** | **Letters** | **Mean** |
| 20 µM CuCl_2_ vs. 20 µM ZnCl_2_ | **0.0003** | 2.5 µM PTO | A | 1.139288 |
| 2 mM CaCl_2_ vs. 20 µM ZnCl_2_ | **0.0017** | 20 µM CuCl_2_ | A | 1.025192 |
| 100 µM CuCl_2_ vs. 20 µM ZnCl_2_ | **0.0026** | 2 mM CaCl_2_ | A | 1.009978 |
| 100 µM CuCl_2_ vs. 2.5 µM PTO | 0.0556 | 100 µM CuCl_2_ | A | 0.994818 |
| 2 mM CaCl_2_ vs. 2.5 µM PTO | 0.1365 | 20 µM ZnCl_2_ | B | 0.805633 |
| 20 µM CuCl_2_ vs. 2.5 µM PTO | 0.2036 |  |  |  |
| 20 µM CuCl_2_ vs. 100 µM CuCl_2_ | 0.9794 |  |  |  |
| 2 mM CaCl_2_ vs. 20 µM CuCl_2_ | 0.9987 |  |  |  |
| 2 mM CaCl_2_ vs. 100 µM CuCl_2_ | 0.9988 |  |  |  |

**Supplementary Table 4: *Statistical analysis of Zn^2+^ effect on MAP decoration of microtubules.*** Listed below are the *p*-values and associated connecting letters report from the post-hoc Tukey HSD test between MAP1b, MAP2C, MAP4, MAP7, MAP9, p150glued, DCX, and Tau. Bolded values indicate statistically significant *p*-values (below α = 0.05). Related to Figure 5H.

| Comparison | *p* | Connecting Letters Report | | |
| --- | --- | --- | --- | --- |
| MAP1b vs. DCX | **<.0001** | **Group** | **Letters** | **Mean** |
| MAP4 vs. DCX | **<.0001** | MAP1b | A | 1.121222 |
| MAP9 vs. DCX | **<.0001** | MAP4 | A | 1.111684 |
| p150glued vs. DCX | **<.0001** | MAP9 | A | 1.081016 |
| MAP7 vs. DCX | **<.0001** | p150glued | A | 1.05963 |
| MAP1b vs. Tau | **<.0001** | MAP7 | A | 1.042955 |
| MAP4 vs. Tau | **<.0001** | MAP2 | B | 0.868078 |
| MAP9 vs. Tau | **<.0001** | Tau | C | 0.575137 |
| MAP2C vs. DCX | **<.0001** | DCX | D | 0.381278 |
| p150glued vs. Tau | **<.0001** |  |  |  |
| MAP7 vs. Tau | **<.0001** |  |  |  |
| MAP2C vs. Tau | **<.0001** |  |  |  |
| MAP1b vs. MAP2C | **0.0001** |  |  |  |
| MAP4 vs. MAP2C | **0.0002** |  |  |  |
| MAP9 vs. MAP2C | **0.0012** |  |  |  |
| Tau vs. DCX | **0.0100** |  |  |  |
| p150glued vs. MAP2C | **0.0055** |  |  |  |
| MAP7 vs. MAP2C | **0.0140** |  |  |  |
| MAP1b vs. MAP7 | 0.8841 |  |  |  |
| MAP4 vs. MAP7 | 0.9339 |  |  |  |
| MAP1b vs. p150glued | 0.9677 |  |  |  |
| MAP4 vs. p150glued | 0.9865 |  |  |  |
| MAP1b vs. MAP9 | 0.9975 |  |  |  |
| MAP9 vs. MAP7 | 0.9975 |  |  |  |
| MAP4 vs. MAP9 | 0.9995 |  |  |  |
| MAP9 vs. p150glued | 0.9999 |  |  |  |
| p150glued vs. MAP7 | 1 |  |  |  |
| MAP1b vs. MAP4 | 1 |  |  |  |

**Supplemental Figures**

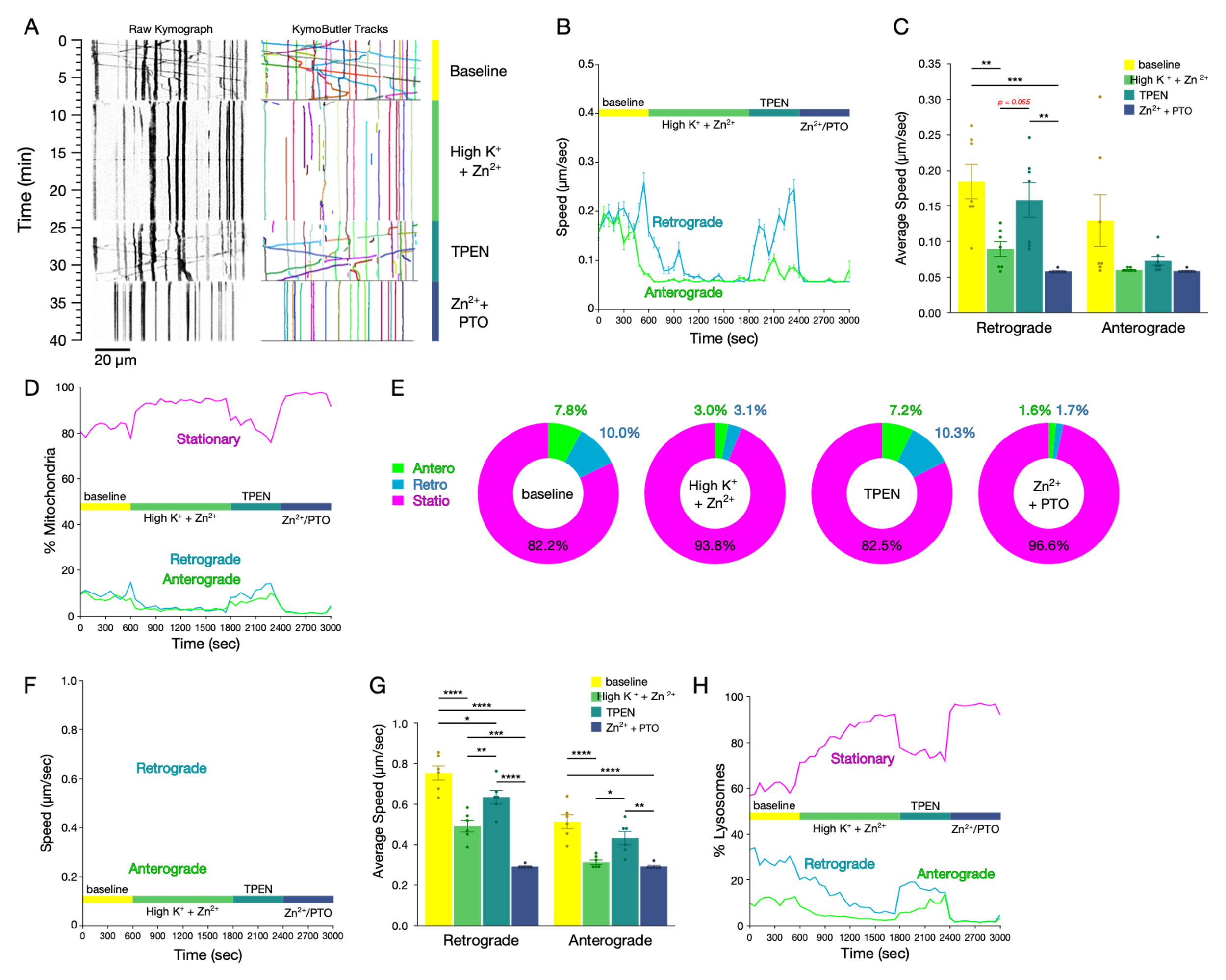

**Figure S1: *Zn^2+^ influx arrests axonal transport of mitochondria and lysosomes.***

(A) Representative kymographs (left) and measured tracks (identified using KymoButler; right) of mito-mCherry motility along axons from primary cultured rat hippocampal neurons prior to (“baseline”; top) or after initiation of Zn^2+^ influx by high K^+^ depolarization (middle-top), following washout and addition of 100 µM TPEN (middle-bottom), and then washout and addition of 100 µM Zn^2+^ and 2.5 µM PTO (bottom). (B) Mean instantaneous speed (±S.E.M.) of mitochondria moving retrograde (blue) or anterograde (green) across baseline, Zn^2+^ influx by high K^+^ depolarization, TPEN, and Zn^2+^ + PTO treatment (60-second binned, representing 7 axons). (C) Mean speed (±S.E.M.) of mitochondria moving in the indicated directions for each condition (n = 7 axons from 3 independent replicates). (D) Proportions of mitochondria moving in the indicated directions across baseline, depolarization, TPEN, and Zn^2+^ + PTO treatment (60-second binned, representing 7 axons). (E) Mean proportions of mitochondrial motility for each condition. (F) Mean instantaneous speed (±S.E.M.) of lysosomes moving retrograde (blue) or anterograde (green) across baseline, Zn^2+^ influx by high K^+^ depolarization, following washout and addition of 100 µM TPEN, and then washout and addition of 100 µM Zn^2+^ and 2.5 µM PTO (bottom). (60-second binned, representing 6 axons). (G) Mean speed (±S.E.M.) of lysosomes moving in the indicated directions for each condition (n = 6 axons from 3 independent replicates). (H) Proportions of lysosomes moving in the indicated directions across baseline, Zn^2+^ influx by high K^+^ depolarization, TPEN, and Zn^2+^ + PTO treatment (60-second binned, representing 6 axons). All experiments were performed in the absence of extracellular Ca^2+^. **** *p* < 0.0001, *** *p* < 0.001, ** *p* < 0.01, * *p* < 0.05, n.s. not significant.

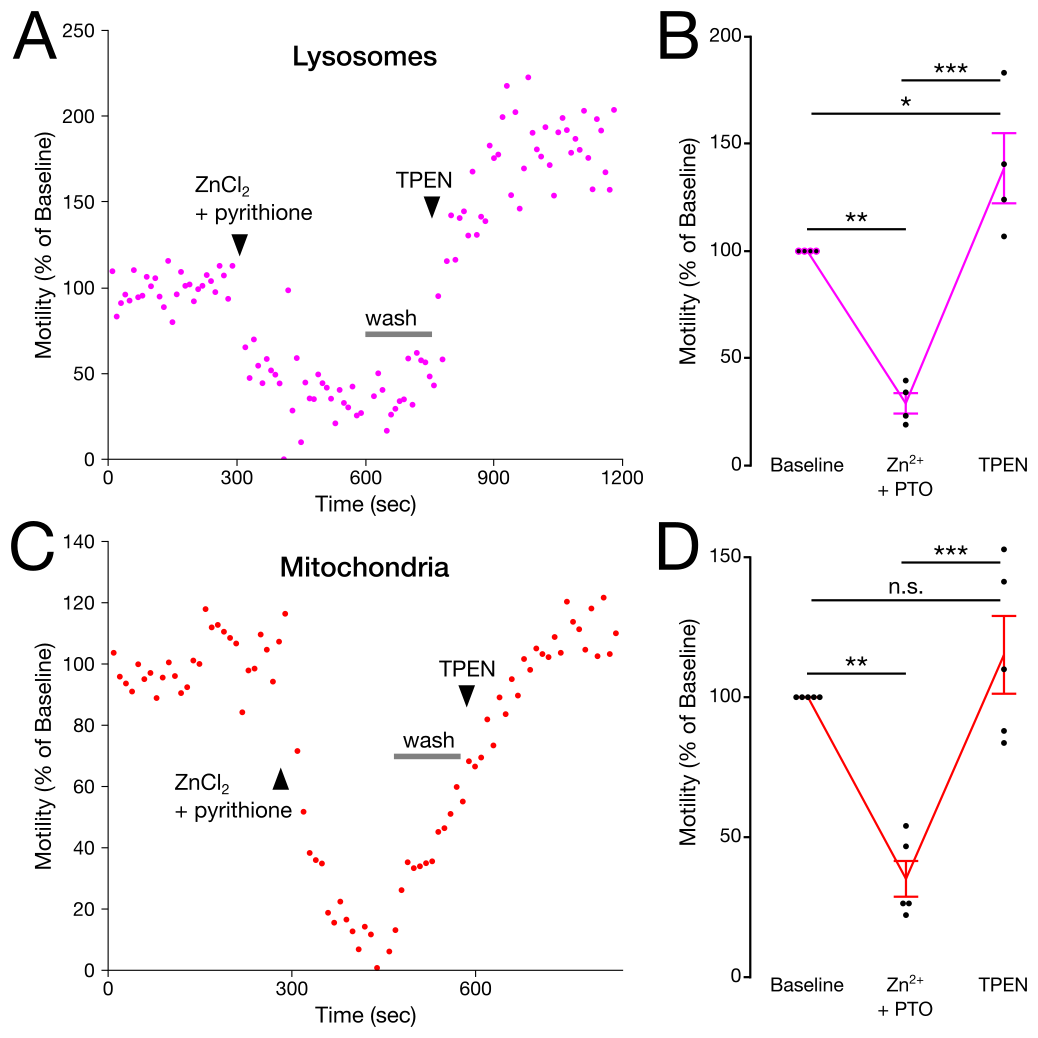

**Figure S2: *Zn^2+^ influx inhibits motility of lysosomes and mitochondria in HeLa cells.*** (A and C) Representative lysosomal (A) and mitochondrial (C) motility (determined using the Total Motility plugin for Fiji; see Methods) in HeLa cells treated with 20 µM ZnCl_2_ and 1.25 µM pyrithione (PTO), then washed and treated with 100 µM TPEN (treatments added at time points indicated on plots). (B and D) Mean (±S.E.M.) lysosomal (B) and mitochondrial (D) motility normalized to their respective baselines (n = 4 and 5 cells, from 4 and 3 independent replicates, respectively). One-way repeated measures ANOVA, with post-hoc Tukey HSD. All experiments were performed in the absence of extracellular Ca^2+^. *** *p* < 0.0001, ** *p* < 0.01, * *p* < 0.05, n.s. not significant.

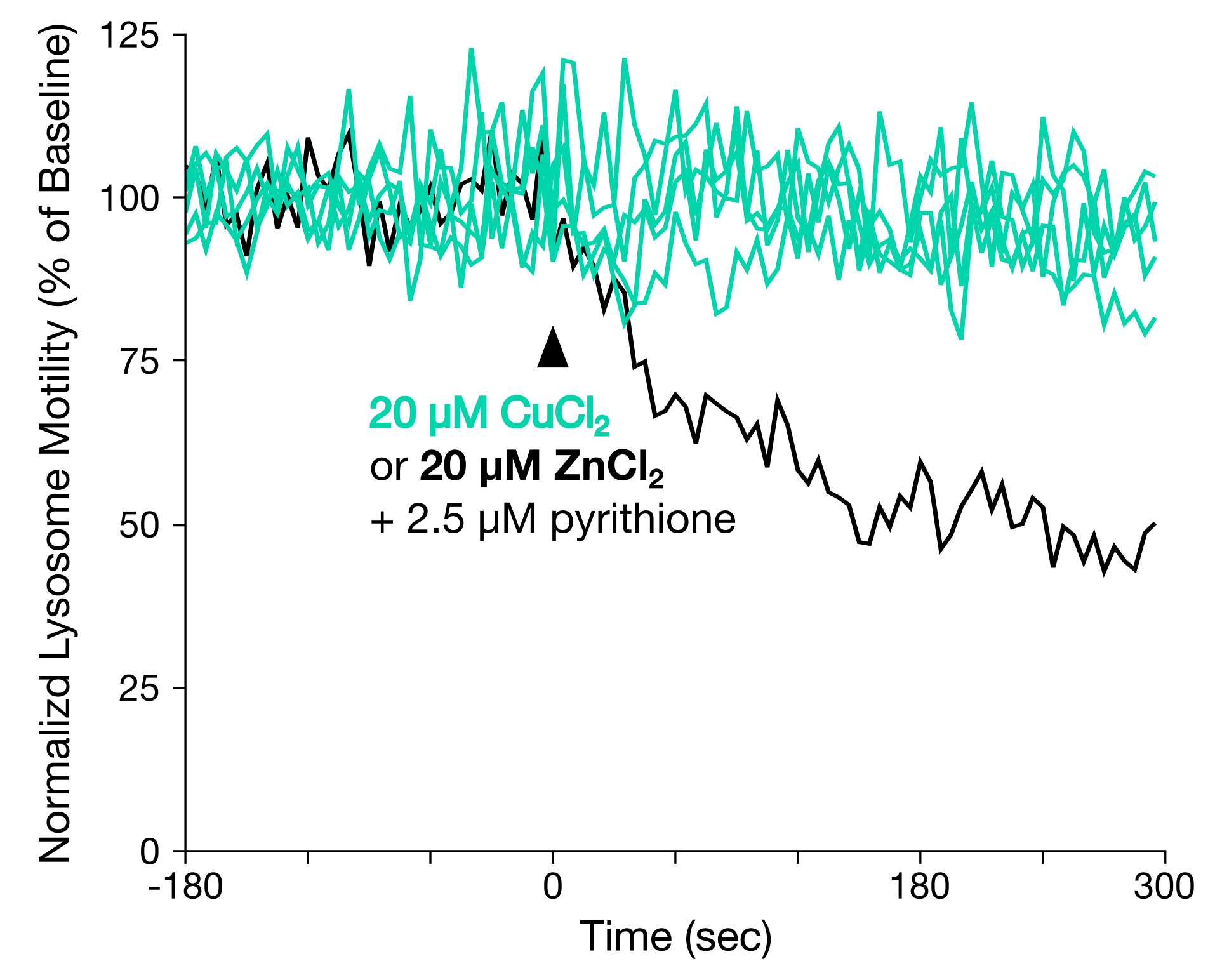

**Figure S3: *Cu^2+^ has no effect on lysosomal motility in HeLa cells.*** (A) Representative traces of lysosomal motility (determined using the Total Motility plugin for Fiji; see Methods) in HeLa cells stained with LysoTracker Red and treated with 2.5 µM pyrithione and either 20 µM CuCl_2_ (teal lines) or 20 µM ZnCl_2_ (black line) at 0 sec. All experiments were performed in the absence of extracellular Ca^2+^.

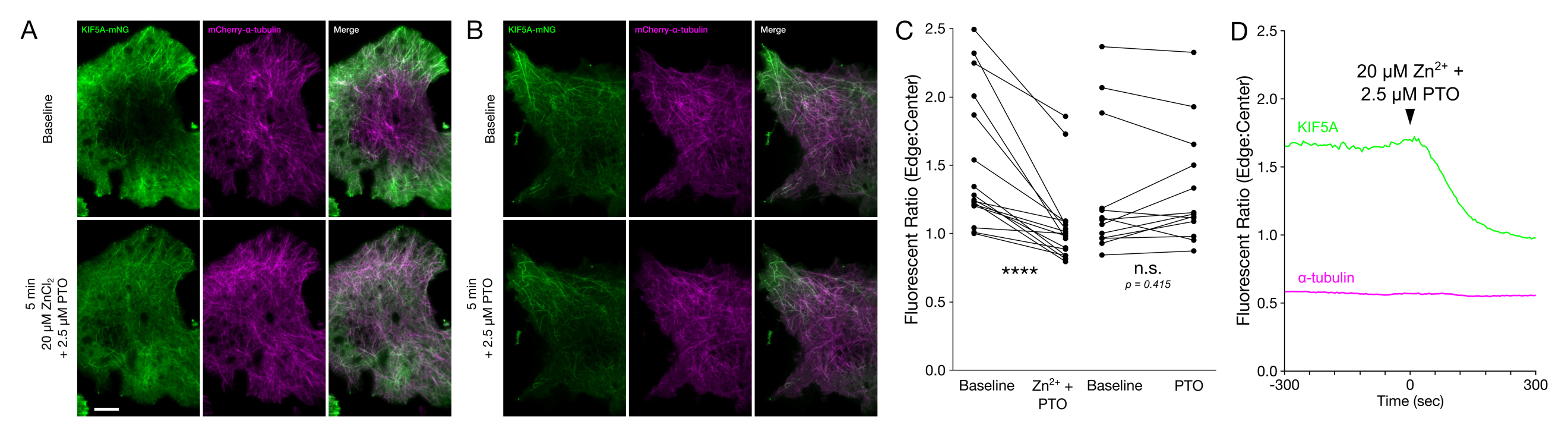

**Figure S4: *Zn^2+^ can bind microtubules and inhibits motor protein activity in situ and in vitro.*** (A-B) Representative oblique illumination micrographs of COS-7 cells expressing KIF5A(1-560)-mNG-FRB (green, left) and mCherry-α-tubulin (magenta, middle), and merged channels (right). Cells were imaged at baseline (top) and 5 minutes after treatment (bottom) with (A) 20 µM ZnCl_2_ and 2.5 µM pyrithione (PTO), or (B) 2.5 µM PTO alone. (C) Ratio of edge-to-center fluorescence of KIF5A(1-560)-mNG -FRB in COS-7 cells as treated in (A) and (B) (Zn^2+^-treated, n = 16 cells from 3 individual biological replicates; PTO control, n = 13 cells from 3 individual biological replicates). Two-tailed paired t-tests. (D) Representative timelapse of edge-to-center fluorescence of KIF5A(1-560)-mNG-FRB (green) and mCherry-α-tubulin (magenta) in COS-7 cells, treated as in (A). All *in situ* experiments were performed in the absence of extracellular Ca^2+^. **** *p* < 0.0001, n.s. not significant.

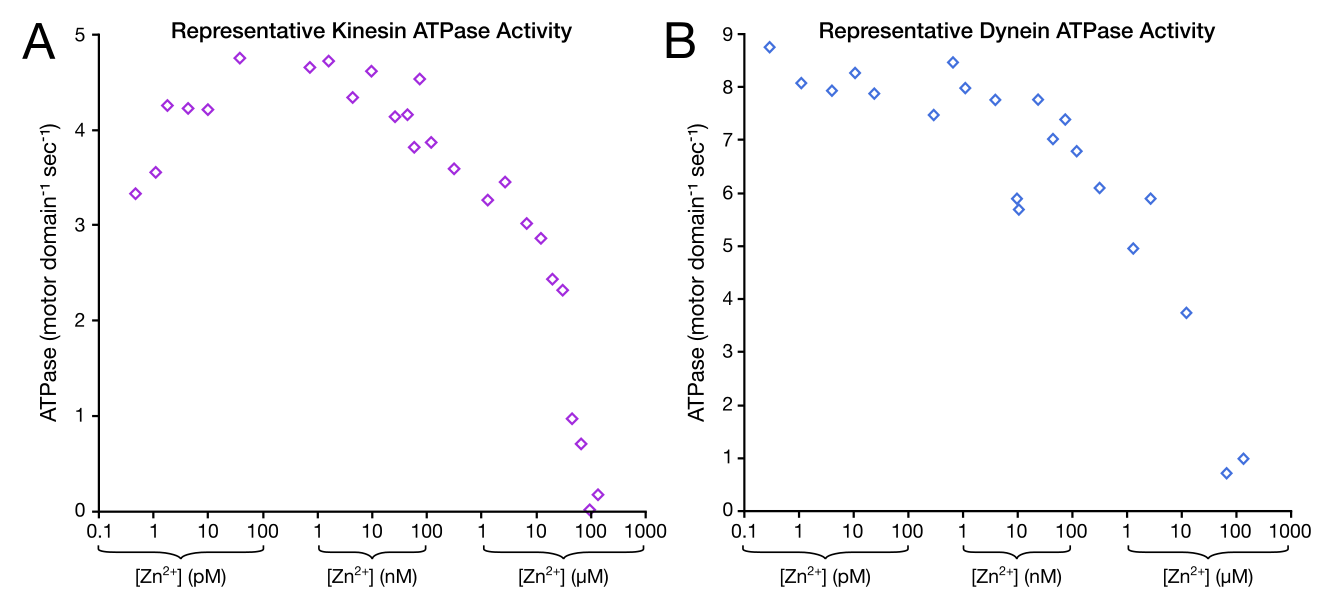

**Figure S5: *Supplemental in vitro analysis of the effect of Zn^2+^* *on kinesin and dynein motor activity.*** (A-B) Representative microtubule-stimulated ATPase activity showing non-normalized ATPase activity per motor domain per second for purified (A) recombinant human kinesin (KIF5A) or (B) a minimally processive, artificially dimerized yeast dynein fragment (GST-dynein_331_) across a range of Zn^2+^ concentrations (from same dataset as Figure 4A-B).

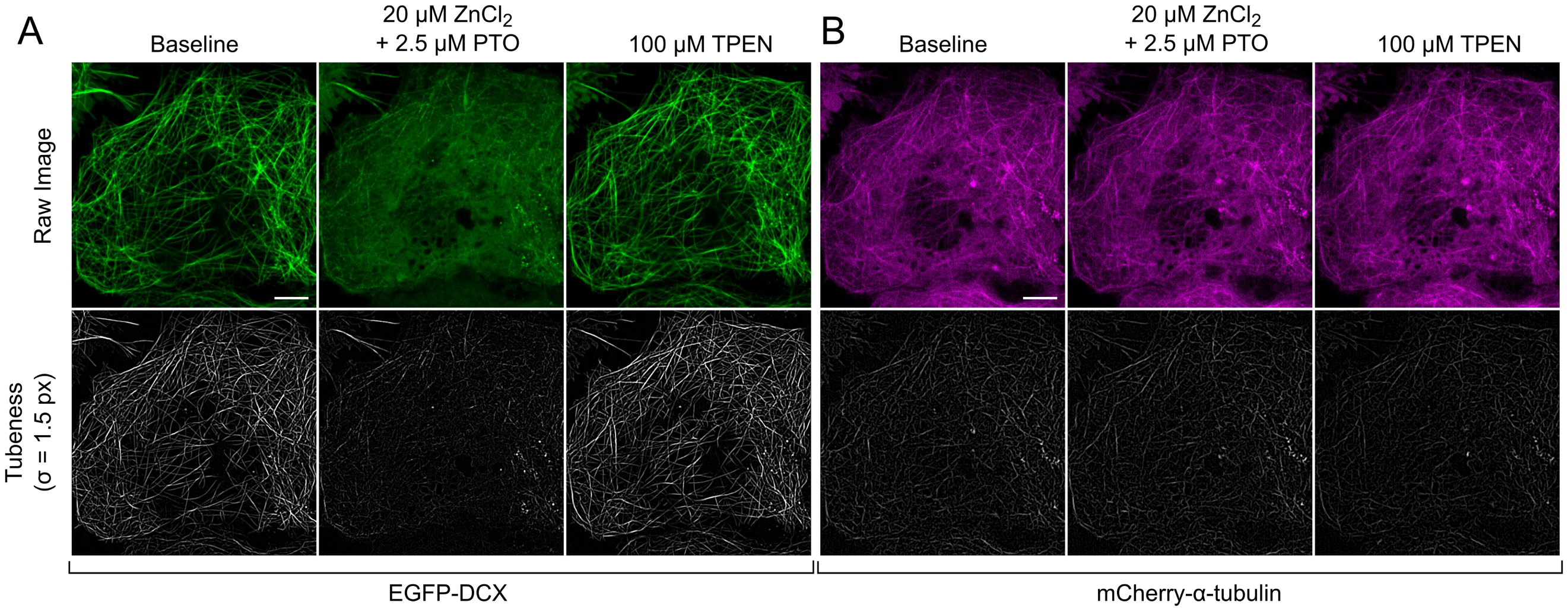

**Figure S6: *Tubeness analysis of EGFP-DCX expressing COS-7 cells with Zn^2+^ and TPEN treatment.*** (A-B) Representative micrographs of COS-7 cells expressing (A) EGFP-DCX and (B) mCherry-α-tubulin before treatment (“baseline”, left), after 5-minute treatment with 20 µM ZnCl_2_ and 2.5 µM PTO (center), or after subsequent treatment with 100 µM TPEN (right). Scale bar = 10 µm. Raw fluorescence images (top) and “Tubeness” analysis (Fiji Plugin) with a sigma, σ, of 1.5 pixels (bottom) are shown.
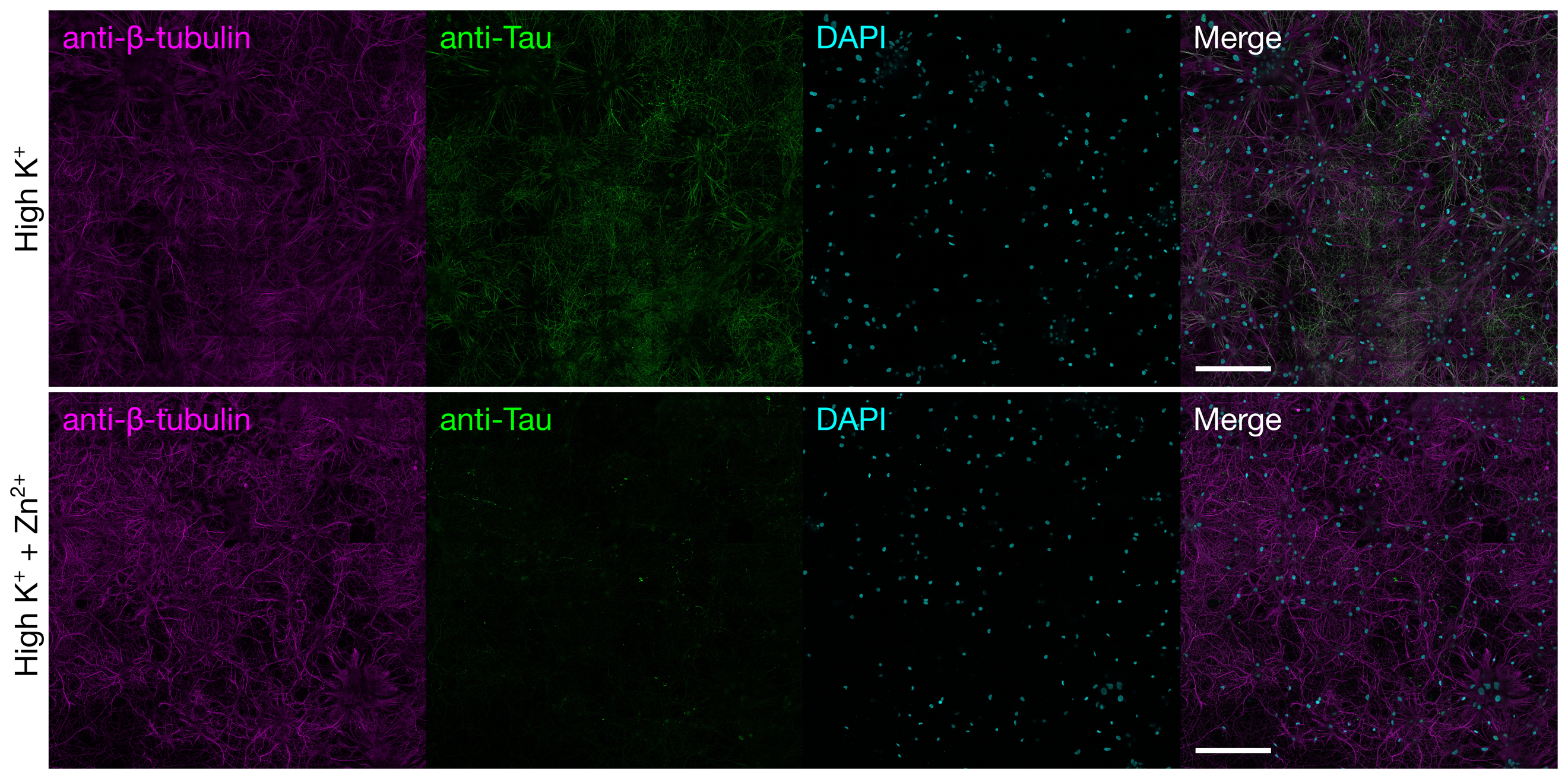

**Figure S7: *Depolarization induced Zn^2+^ influx promotes detachment of endogenous neuronal Tau in situ.*** Representative immunofluorescence micrographs (15 x 15 stitched image grid, 5% overlap) of methanol-fixed primary rat hippocampal neurons depolarized with 50 mM KCl in the absence (top) or presence (bottom) of 100 µM ZnCl_2_. Fixed cells were incubated with anti-β-tubulin monoclonal antibodies (magenta, left), anti-Tau polyclonal antibodies (green, left center), DAPI (cyan, right center), and channels are shown merged (right). Scale bar = 200 µm. All experiments were performed in the absence of extracellular Ca^2+^.

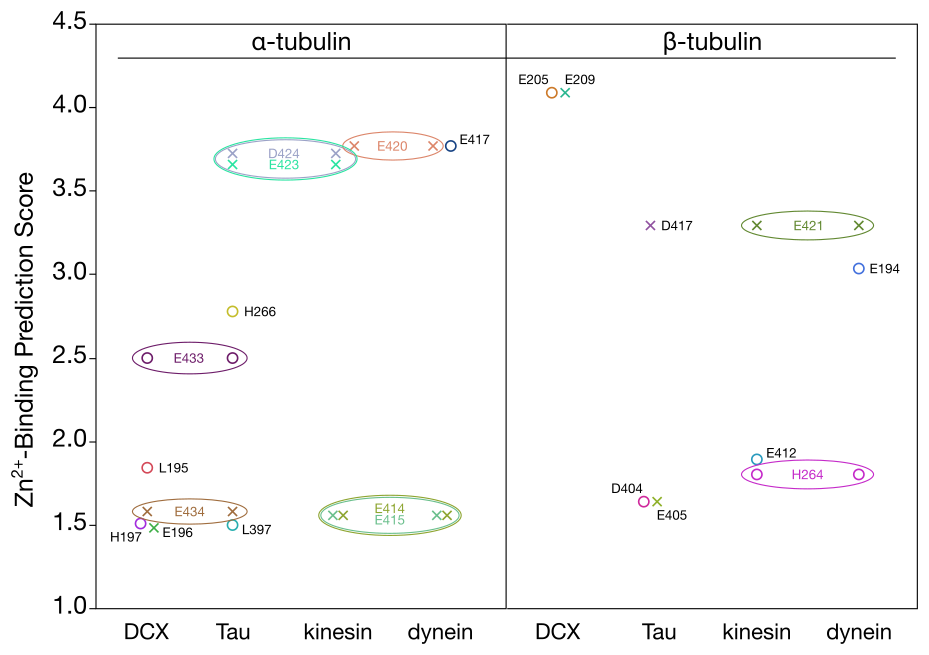
**Figure S8: *Predicted Zn^2+^ binding near MAP binding sites.*** Zn^2+^-binding prediction scores of α-tubulin and β-tubulin amino acid residues that directly interact with the indicated MAPs (“X” markers) and residues within 3Å of MAP-interacting residues (open circle markers). Residues that are shared between MAPs are represented by enclosed ovals.

**Supplemental Video Captions**

**Video S1: *Depolarization-induced influx of Zn^2+^ arrests axonal transport of lysosomes.*** LAMP1-mCherry labeled lysosomes moving along axons of primary rat hippocampal neurons during baseline, depolarization induced Zn^2+^ influx (High K + Zn^2+^), and washout followed by TPEN treatment. Related to Figure 1.

**Video S2: *Depolarization-induced influx of Ca^2+^does not inhibit axonal transport of lysosomes.*** LAMP1-mCherry labeled lysosomes moving along axons of primary rat hippocampal neurons during baseline, depolarization induced Ca^2+^ influx (High K + Ca^2+^), and washout. Related to Figure 1.

**Video S3: *Zn^2+^ influx arrests axonal transport of mitochondria.*** mito-mCherry labeled mitochondria moving along axons of primary rat hippocampal neurons during baseline, depolarization induced Zn^2+^ influx (High K + Zn^2+^), washout followed by TPEN treatment, and washout followed by Zn^2+^ and pyrithione (PTO) treatment. Related to Figure S1.

**Video S4: *Zn^2+^ inhibits organellar motility with nanomolar IC_50_* *in HeLa cells*.** Simultaneous imaging of Zn^2+^ (using GZnP2 sensor) and either lysosome (LAMP1-mCherry) or mitochondria (mito-mCherry) motility in HeLa cells. Related to Figure 2 and Figure S2.

**Video S5: *Zn^2+^ inhibits KIF5A movement in situ in a dose dependent manner***. Peroxisome dispersion assays in COS-7 cells expressing KIF5A(1-560)-mNG-FRB and PEX-mRFP-2xFKBP, during baseline, treatment with pyrithione (PTO) and varying concentrations of Zn^2+^, and subsequent treatment with Zotarolimus. Related to Figure 3.

**Video S6:** ***Zn^2+^ redistributes KIF5A in COS-7 cells.*** Distribution of KIF5A(1-560)-mNG-FRB and mCherry-α-tubulin in COS-7 cells during baseline and after Zn^2+^ and pyrithione (PTO) treatment. Related to Figure S4.

**Video S7:** ***Zn^2+^ promotes the dissociation of doublecortin from microtubules in situ.*** Microtubule decoration of EGFP-DCX in COS-7 cells during baseline, Zn^2+^ and pyrithione (PTO) treatment, and Zn^2+^ chelation with 100 µM TPEN. Related to Figure 5.
